# Day-night and seasonal variation of human gene expression across tissues

**DOI:** 10.1101/2021.02.28.433266

**Authors:** Valentin Wucher, Reza Sodaei, Raziel Amador, Manuel Irimia, Roderic Guigó

## Abstract

Circadian and circannual cycles trigger physiological changes whose reflection on human transcriptomes remains largely uncharted. We used the time and season of death of 932 individuals from GTEx to jointly investigate transcriptomic changes associated with those cycles across multiple tissues. Overall, most variation across tissues during day-night and among seasons was unique to each cycle. Although all tissues remodeled their transcriptomes, brain and gonadal tissues exhibited the highest seasonality, whereas those in the thoracic cavity showed stronger day-night regulation. Core clock genes displayed marked day-night differences across multiple tissues, which were largely conserved in baboon and mouse, but adapted to their nocturnal or diurnal habits. Seasonal variation of expression affected multiple pathways and it was enriched among genes associated with the immune response, consistent with the seasonality of viral infections. Furthermore, they unveiled cytoarchitectural changes in brain regions. Altogether, our results provide the first combined atlas of how transcriptomes from human tissues adapt to major cycling environmental conditions.

## Introduction

The yearly and daily movement of the earth around the sun and around itself has created a continuously changing environment since the origin of life to which all organisms across the phylogenetic spectrum have adapted. In mammals, in particular, behavioral adaptations to daily rhythm include the regulation of sleep and feeding cycles. Recent studies have investigated how the physiological responses to the daily cycle are reflected at the transcriptomic level. These studies have reported large-scale circadian gene expression oscillations in mice (Zhang et al. 2014), baboons (Mure et al. 2018), and humans (Ruben et al. 2018; Anafi et al. 2017). Unlike the circadian process, circannual rhythm in mammals has been less studied. Animals exhibit a range of behavioral and physiological adaptations in different seasons, such as hibernation and alterations of the coat color in polar animals (Drew et al. 2007; Ferreira et al. 2017). One way in which mammals regulate their seasonal reproductive behavior, growth, food intake, and migratory behavior is via the brain-gonadal and other hormonal axes (Hanon et al. 2008; Dardente et al. 2014; Lomet et al. 2018). In humans, numerous pathologies present a strong seasonal pattern, which is particularly prominent for many infectious diseases, but also observed in complex cardiovascular and psychiatric disorders (Watson et al. 1984; Philpot et al. 1989; Pell et al. 1999; Pell and Cobbe 1999; Torrey et al. 2000; Owens and McGorry 2003; Xu et al. 2013; Li et al. 2017a; Sharon et al. 2019). Nevertheless, despite its relevance for human physiology and disease, genome-wide studies on circannual rhythms are scarce. Castro Dopico *et al*. (2015) analyzed the transcriptome of white blood cells from children in Germany, from individuals affected with type 1 diabetes from the UK, and from asthmatic cohorts from distinct geographical locations (Dopico et al. 2015). They found many genes with seasonal expression profiles, inverted between Europe and Oceania. These seasonal expression profiles were prominent in genes from the immune system. To our knowledge, however, there are no genome-wide studies of the transcriptional impact of these adaptations to seasonal variation across multiple human tissues.

Here, we leverage on the deep transcriptome data across human tissues produced by the GTEx consortium (16,151 RNA-seq samples of 932 post-mortem human donors from 46 tissues) (GTEx Consortium 2020) to investigate the transcriptional impact of circadian and circannual rhythms in an unprecedented number of tissues. GTEx transcriptional measurements are taken exclusively at the donor’s death; therefore, there is a single time-point measure per individual. In addition, GTEx metadata only includes the time of the day and the season of death, but not the actual day, the week or even the month of death. This prompted us to artificially discretize circadian and circannual variation into day-night and season-specific variation. Despite all these caveats, we show that, when aggregated over many individuals, these transcriptional snapshots randomly distributed along time create temporal trajectories that recapitulate day-night and seasonal transcriptional variation, and they constitute, therefore, a unique resource to investigate this variation.

## Results

### Tissue-dependent day-night variation in gene expression

We first used MetaCycle (Wu et al. 2016), a suite designed to analyze rhythmic data, and that has been previously used to investigate circadian patterns (e.g. Ruberto *et al*. (Ruberto et al. 2021) and Mishra *et al*. (Mishra et al. 2021)). While MetaCycle uses three methods to identify oscillating genes, only Lomb-Scargle is suited for GTEx data, due to the uneven distribution of the time of death. Using this method, from the 18,018 protein-coding genes expressed in at least one tissue (median TPM ≥ 1), we identified 187 (1%) that were circadian in at least one tissue (non-adjusted *P* ≤ 0.05; Table S1). This number is much smaller than that previously reported in baboon (Mure et al. 2018), where 82% of the protein-coding genes had circadian patterns in at least one of the 64 studied tissues. Lack of power to detect genes with circadian gene expression patterns can be partially attributed to the characteristics of the GTEx resource. As reported, the GTEx metadata only includes the time of the day and the season of death, and there is no information about the location of death. Therefore, we focused only on the individuals in which the time of death could be unequivocally classified as either day [8:00-17:00) (351 individuals) or night [21:00-5:00) (315 individuals), and excluded those in which death occurred outside these intervals (the twilight zone; 222 individuals) (Fig. 1A). We then identified genes that were differentially expressed between day and night.

**Fig. 1:**
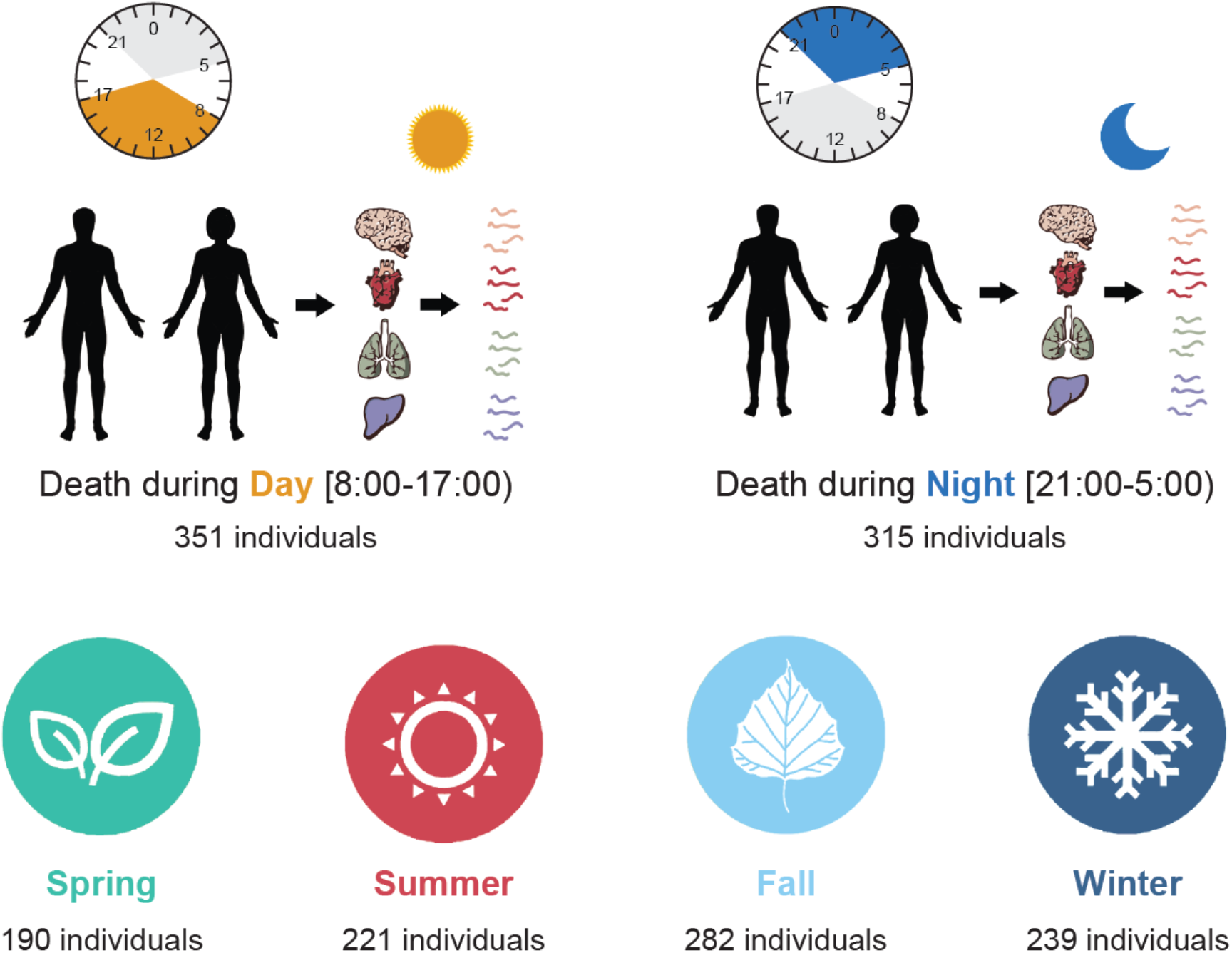
Day-night and seasonal classification of the GTEx samples. (**A**) Samples from GTEx were classified as either day or night depending on the time of death of the donor, respectively between 08:00 and 17:00 (351 individuals) and between 21:00 and 05:00 (315 individuals). (**B**) Samples from GTEx were assigned to the different seasons (spring, summer, fall and winter), according to the reported season of death of the donor.

For this purpose, we carried out a differential gene expression analysis using *voom/limma* (Law et al. 2014) controlling for the effects of season of death, sex, body mass index, age, and post-mortem interval. We performed an initial comparison across tissues with high sensitivity, applying loose cut-offs of a non-adjusted *P* ≤ 0.05 and an absolute log_2_ fold-change ≥ 0.1. From the 18,022 protein-coding genes expressed in at least one tissue (median TPM ≥ 1), we found that 12,530 (70%) were differentially expressed between day and night in at least one of the 46 tested tissues (day-night genes, Supplementary Dataset 1), which is a number in line with the results found in baboon. However, it should be noted that, while this approach increases the power to detect significant changes in gene expression, it does not necessarily lead to the identification of all genes with circadian patterns of gene expression, since the circadian peaks may occur at the twilight time zones that we are ignoring. Consistently, most of the gene-tissue pairs detected by MetaCycle as circadian were also identified as day-night by our analyses (210/339, 62%; Table S1), with most exceptions peaking at or around the twilight (Table S2).

Per tissue, 5.5% of expressed protein-coding genes were day-night on average. The tissue with the largest number of day-night genes was lung, with 2,418 genes (17.2% of genes expressed in this tissue; Fig. 2A). Other tissues with a proportionally large number of day-night genes were the heart left ventricle (2,202 genes, 19.2%) and whole blood (1,900 genes, 19%). On the other hand, the tissue with the fewest differentially expressed day-night genes was salivary gland, with only 85 genes (0.63% of genes expressed in this tissue; Fig. 2A). Other tissues with proportionally low number of day-night genes were colon transverse (92, 0.67%) and testis (105, 0.66%). Brain regions also showed a relatively small number of day-night changes (ranging from 0.86% to 7.8%; Fig. 2A). Caudate was the brain region with the highest number of day-night genes (1,026 7.8%), followed by the cerebellum (766, 5.7%), which was reported to have a sleep stage–dependent activity (Canto et al. 2017). Some tissues showed a bias in the number of genes overexpressed during the day (diurnal genes) versus the night (nocturnal genes) (Fig. 2A and Table S3). Stomach was the tissue with the strongest diurnal preference, while non-sun exposed skin was the tissue with the strongest nocturnal preference. In contrast, sun exposed skin showed diurnal preference (0.7 log_2_ day/night genes ratio), suggesting that UV plays a role in the activation of gene expression.

**Fig. 2:**
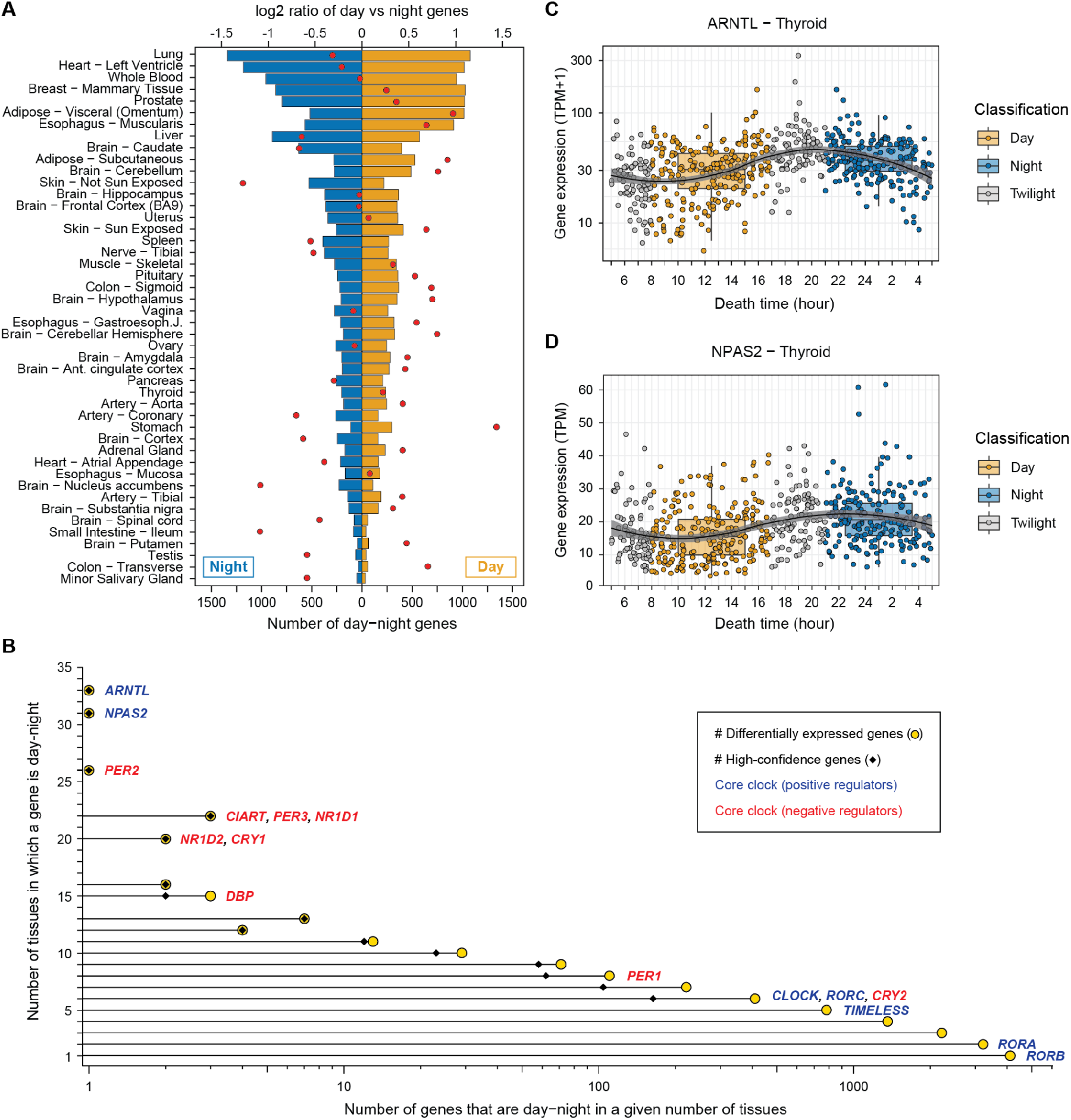
Distribution of the day-night genes in the GTEx tissues. (**A**) Number of genes found as day-night, i.e. genes differentially expressed between day and night (barplots; bottom x-axis), and log_2_ ratio between the number of genes up-regulated during the day vs the night (red dots; top x-axis) for each tissue. Tissues are sorted by the total number of day-night genes. (**B**) Yellow dots represent the number of tissues in which a given gene is classified as day-night (y-axis) vs the number of genes that are classified as day-night in that number of tissues (x-axis, log_*10*_ scale). Gene names for core clock genes are shown next to the dot with the corresponding number of tissues in which they are identified as day-night. Black diamonds show the number of high-confidence day-night genes per number of tissues (see Fig. S3 for further details). (**C**) Expression of *ARNTL* in the thyroid in GTEx samples at the time of death of the GTEx donors (in hours). The colors of the dots represent the classification of the individuals according to their time of death: during the day (yellow), during the night (blue), or during twilight (grey). The samples classified as twilight were discarded for the day-night analysis. The “circadian” curve was created using the *geom_smooth* function from *ggplot2* in R with the ‘loess’ method.

### Recurrent day-night variation in gene expression across tissues

We next computed the number of tissues in which a given gene showed a day-night pattern (Fig. 2B). We found that, on average, day-night genes exhibited a day-night pattern in 2.6 tissues out of 39 tissues in which they had detectable expression (median TPM ≥ 1), indicating a rather tissue-specific response to day-night cycles. Overall, we found that genes identified as day-night in more than one tissue significantly tended to be consistently up-regulated in either day or night, and this consistency largely increased with the number of tissues in which genes were detected as day-night (Table S4; see Methods).

We focused on a set of 16 genes that form the molecular core clock and are the main regulators of the circadian rhythm (Bargiello et al. 1984; Shearman et al. 2000; Liu et al. 2008) (Table S5). With the exception of *RORC* and *RORB*, the core clock genes were expressed in almost all tissues, and, with the exception of *RORB*, they showed differential day-night expression in multiple tissues (Fig. 2B). This is in agreement with the hypothesis that one molecular clock program is present in all tissues but the circadian processes triggered downstream are highly tissue-specific (Scheiermann et al. 2013). Core clock genes usually have effect sizes (difference between day and night expression) that are larger than those of non-core clock day-night genes (Fig. S1). *ARNTL*, a positive regulator of the core clock, was the gene with a day-night pattern in the largest number of tissues (33 tissues; Fig. 2B). Its expression at the time of death, aggregated across the GTEx donors, nicely captures the known circadian behaviour of this gene (Fig. 2C), strongly indicating that, in spite of all the caveats associated with the data collection and available metadata, the GTEx data can be effectively used to investigate gene expression patterns during the day-night cycle. A recent study described *ARNTL* as the main regulator of the inter-tissue timekeeping function in mouse (Welz et al. 2019), and our results suggest that it may play a similar role in humans. Similarly, *NPAS2* was the second gene with a day-night pattern in the largest number of tissues (31 tissues; Fig. 2B,D) and has also been reported as being circadian in many mouse (Li et al. 2018) and baboon (Mure et al. 2018) tissues.

Among the core clock genes, the thyroid had the highest number of day-night cases (12 out of 16 genes; Fig. 3A). In contrast, we did not detect any of them as day-night in stomach, testis, and vagina, consistent with their overall low number of day-night changes (Fig. 2A), and with previous studies in testis in rodents (Bittman 2016). Clock genes showed a largely consistent pattern of day-night expression across tissues (i.e. they were either consistently up- or down-regulated at the same time of the day; Fig. 3A). One exception was *NR1D2*, whose expression was consistently higher diurnally in brain subregions but nocturnally in all other tissues (Fig. 3B and Fig. S2). For the core clock genes that showed day-night differences in both human and baboon in any of the available 20 homologous tissues (Table S6), we found that most orthologs had a similar behavior in both species (59 similar vs. 15 opposite gene-tissue pairs; *P* = 1.277 × 10^-7^ one-sided binomial test; Fig. 3C), consistent with their shared diurnal regimes. In contrast, core clock genes in mouse (a preferentially nocturnal mammal) largely showed the opposite behavior than their human counterparts (13 similar vs. 38 opposite gene-tissue pairs; *P* = 3.105 × 10^-4^ one-sided binomial test; Fig. 3C and Table S7).

**Fig. 3:**
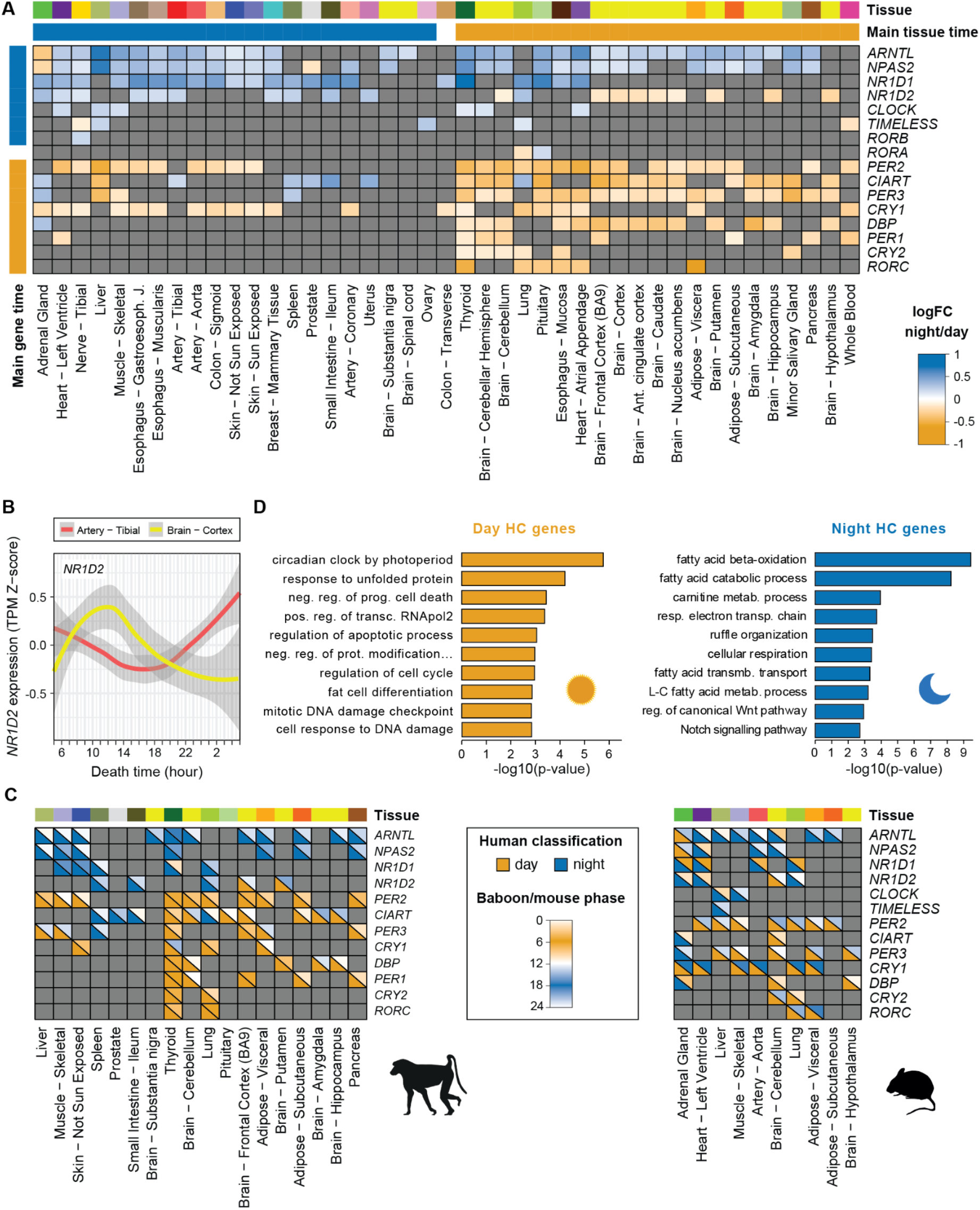
Day/night differential expression of the core clock genes in human, baboon, and mouse. (**A**) Day/night differential expression of the 16 core clock genes in GTEx tissues. Cells in the matrix are colored according to the log_2_ fold-change obtained with the *voom-limma* pipeline, from yellow (day) to blue (night). Genes without significant effects were colored in grey. In addition, we labelled each gene (“Main gene time”) as day (night) if it was up-regulated in more tissues during the day (night) than during the night (day). We labelled the tissues similarly (“Main tissue time”) depending on the number of genes that were up-regulated during the day (night) on that tissue. One gene, *RORA*, and one tissue, the aorta, were upregulated in the same number of times during the day and the night, and have not been labelled. Genes and tissues have been sorted according to i) their main time and ii) their number of significant effects. (**B**) Expression of *NR1D2* in the artery - tibial (red) and in the brain - cortex (yellow) GTEx samples at the time of death of the GTEx donors (in hours). Curves were obtained from the Z-score of the expression TPMs using the *geom* smooth function from *ggplot2* in R with the ‘loess’ method (**C**) Comparison of human and baboon (left) and human and mouse (right) orthologous core clock genes in common tissues. Only gene-tissue pairs that are significant for both compared species are shown. Cells are separated into two with i) on the bottom left, the day-night classification of the gene in the tissue in human, and ii) on the top right, the baboon/mouse phase obtained from Mure *et al*. (Mure et al. 2018) or Li *et al*. (Li et al. 2018), respectively. (**D**) Top 10 enriched Gene Ontology Biological Process annotations for day and night high-confidence genes separately. See Methods for details.

Next, we identified 445 genes with a consistent day (282) or night (165) pattern in multiple tissues (*P* ≤ 0.05 two-sided binomial test across all tissues, brain regions and/or across non-brain tissues; see Fig. S3, Table S8 and Methods). We defined these genes as “high-confidence day-night genes” (Fig. 2B). High-confidence day-night genes included most clock genes (with the exception of *TIMELESS, RORA* and *RORB*) and were significantly enriched for circadian and photoperiodism related Gene Ontology (GO) terms (Fig. 3D and Table S9). Moreover, this gene set significantly overlapped those annotated as circadian in the Circadian Gene DataBase in humans (Li et al. 2017b) (87 genes, *P* = 3.074 × 10^-8^ one-sided Fisher test), and substantially expands the set of genes known to be varying during the day-night cycle. Additional GO terms were enriched among high-confidence day-night genes, including apoptosis and cell cycle regulation (genes peaking during the day) or fatty acid metabolism, cellular respiration and various signalling pathways (genes peaking during the night)(Fig. 3D and Table S9). An example of high-confidence day-night gene with previously unknown circadian variation is *THRA* (Fig. S4A), a thyroid hormone receptor that peaks at night in 15 tissues (including the thyroid), the expression of which was found disrupted in the hypothalamic structures of rats in constant darkness or lighting (Klimina et al. 2019). Other examples of high-confidence day-night genes present in a large number of tissues are the ribosomal protein *RPS26* (day in 16 tissues), the nuclear pore complex protein *NPIPB5* (day in 16 tissues), and the transcription corepressor *TRIM22* (day in 13 tissues) (Fig. S4B-D).

One of the physiological signatures of the day-night rhythmicity is the sleep-wake cycle. Therefore, we next focused on a set of 254 protein-coding genes expressed in GTEx and that were previously reported to increase the risk of insomnia or to be associated with other sleep traits in humans (Jansen et al. 2019) (Table S10). From this gene set, 186 (73.2%) exhibited a significant day-night variation in at least one tissue, closely matching the genome-wide behavior across tissues (Fig. S5). Moreover, none of them belonged to the core clock and they did not significantly overlapped with our set of high-confidence day-night genes (7 genes in common; *P* = 0.49, one-sided Fisher test) or with those annotated as circadian in the Circadian Gene DataBase in human (Li et al. 2017b) (*P* = 0.31, one-sided Fisher test over expressed genes). These results thus suggest that genes involved in sleep disorders and traits do not appear to be particularly impacted by circadian or day-night gene expression patterns. However, four sleep genes were high-confidence day-night (Fig. S6) and annotated as circadian in the Circadian Gene DataBase (Li et al. 2017b) either in human (*PC*) or in other species (*PITPNC1* and *PDE4B* in mouse and *QSOX2* in *Arabidopsis thaliana*), pointing at a potential relevance, in specific cases, on day-night regulation of sleep behaviours.

### Tissue-dependent seasonal variation in gene expression

Next, we analyzed the seasonal variation of gene expression. To make the analyses consistent with those of day-night patterns and to minimize the impact of GTEx reporting only the season and not the actual day of death (see Methods), we set out to identify season-specific genes, i.e. genes that are differentially highly or lowly expressed in one particular season vs. the others. This minimizes some of the impact of measurements taken around the seasonal boundaries. Including as co-variates day-night variation as well as sex, BMI, age and post-morten interval, and using a comparable definition of differential gene expression (raw *P* ≤ 0.05 and absolute log_2_ fold-change ≥ 0.1, see Methods), we found that 16,408 (91.1%) of all expressed protein-coding genes were differentially expressed in at least one season in at least one tissue (hereafter seasonal genes; Supplemental Dataset 2). There were no large differences in the number of seasonal genes across seasons, ranging from 12,026 genes in summer to 13,192 in fall. Per tissue and per season, the average number of seasonal genes were similar among seasons and comparable to day-night patterns (5.3% in spring, 4.9% in summer, 6.5% in fall, and 5.3% in winter, compared with 5.5% day-night genes), but there were many more unique seasonal genes than day-night genes per tissue when all seasons were considered together (17.7%). The effect sizes of seasonal genes were also similar to those observed in day-night genes (Fig. S7).

In stark contrast to day-night patterns, the tissue with the highest proportion of genes showing seasonal changes was testis (25.6% of expressed genes; Fig. 4A), with the highest variation occurring in fall and spring (Fig. S8). Most other tissues with large seasonal changes were brain subregions in summer, fall, and winter. Most of these tissues did not show a clear bias in the direction of the expression changes (i.e. the number of season-specific up- or down-regulated genes was similar; Fig. S8 and Table S11). Testis, however, exhibited a massive gene up-regulation in fall and down-regulation in spring.

**Fig. 4:**
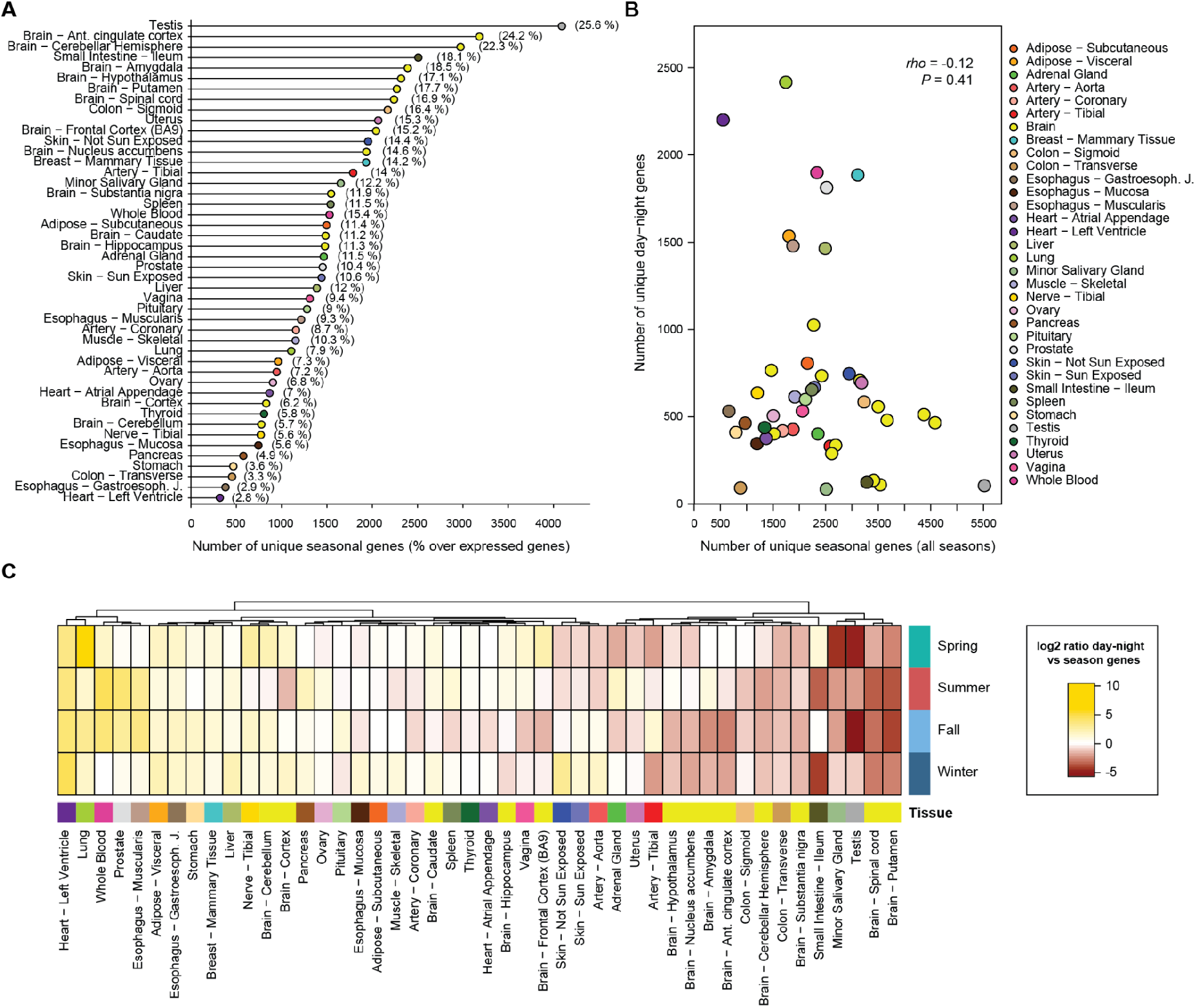
Distribution of the seasonal genes in the GTEx tissues. (**A**) Number of unique genes found as seasonal (x-axis), i.e. genes differentially expressed in at least one season when compared to the others, per tissue (y-axis). The numbers in parentheses represent the percentage of unique seasonal genes in a given tissue over the number of expressed genes in that tissue. (**B**) Number of unique seasonal vs. day-night genes per tissue. Statistics from a Spearman’s correlation are shown. Tissue colors correspond to the GTEx color panel. (**C**) log_2_ ratio between the number of day-night genes vs seasonal genes for each tissue and season separately. The higher the log_2_ value, the more day-night genes compare to the number of seasonal genes. Tissues were clustered using Euclidean distance and the Ward’s method.

Although the overlap between seasonal and day-night genes was statistically significant for most tissues, most differentially regulated genes were unique to one or the other type of variation (Table S12). Moreover, consistent with the distinct set of top varying tissues, the number of day-night and seasonal genes across tissues did not correlate (Spearman’s rho = - 0.12; Fig. 4B). Tissues with larger day-night than seasonal variation included various tissues from the thoracic cavity (e.g. lung and the heart’s left ventricle), which may reflect changes in heart rate and breathing patterns between day and night (Penzel et al. 2003). On the other hand, tissues with more seasonal than day-night genes included most brain subregions and gonadal tissues, likely mirroring the involvement of brain-gonadal axis in regulating seasonal physiology and behavior (Plant 2015) (Fig. 4C).

### Recurrent seasonal variation of gene expression across tissues

On average, seasonal genes showed seasonal expression in a number of tissues comparable to that of day-night genes: 2.5 in spring, 2.5 in summer, 3 in fall, and 2.5 winter, compared with 2.6 tissues for day-night genes. These genes also tended to be consistently up- or down-regulated across tissues. Therefore, as per day-night variation, we defined sets of high-confidence seasonal genes that varied in a consistent manner across multiple tissues for each season separately (Table S13; see Methods). In total, we identified 1,748 unique genes: 308 in spring (138 up and 170 down), 361 in summer (158 up and 203 down), 1,072 in fall (691 up and 381 down), and 322 in winter (89 up and 233 down) (Fig. 5A, Table S13). The top enriched gene functions were largely specific for individual seasons, although some regulatory categories such as transcription and translation were shared by multiple sets (Fig. 5B and Table S14). As expected, multiple immune-related terms were enriched in fall and winter.

**Fig. 5:**
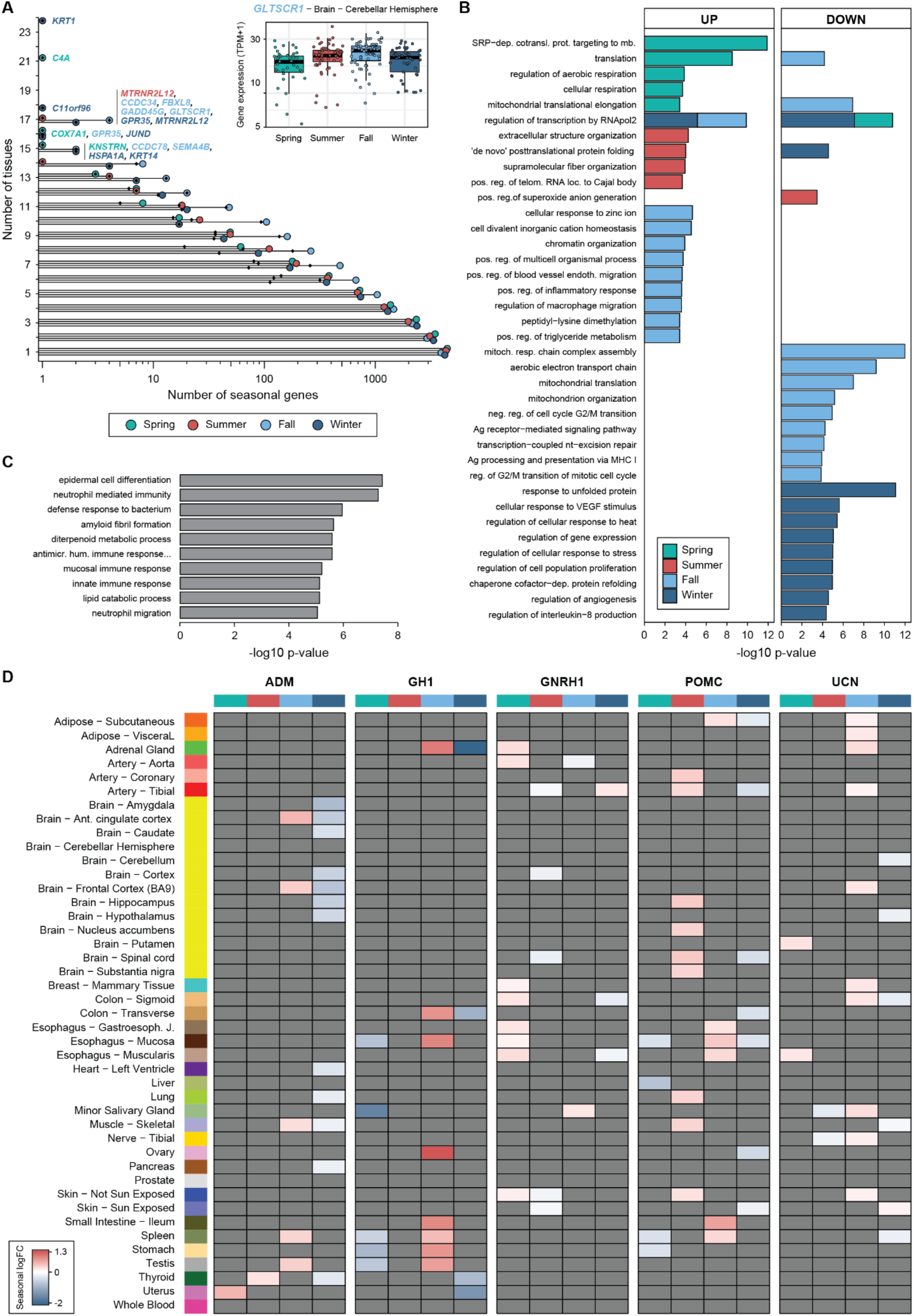
Seasonal genes and associated functions. (**A**) Distribution of the number of tissues (x-axis, log_*10*_ scale) in which genes were classified as seasonal (green: spring, red: summer, light blue: fall, dark blue: winter). The names of the genes found as seasonal in a large number of tissues for a given season are depicted and the font color corresponds to their respective season. Black diamonds show the number of high-confidence seasonal genes per number of tissues. (**B**) Enriched Gene Ontology biological process annotations for up and down high-confidence seasonal genes separately per season (green: spring, red: summer, light blue: fall, dark blue: winter). Top terms with a raw *P* < 0.005 are shown; see Methods for details. (**C**) Gene Ontology enrichment of Biological Processes of the set of 192 strongly seasonal genes as computed by Enrichr (Chen et al. 2013; Kuleshov et al. 2016). (**D**) Seasonal log_2_ fold-change of the five hormone-coding genes included in at least one high-confidence seasonal gene set. Genes without significant effects are colored in grey.

An interesting example of high-confidence seasonal gene is *GLTSCR1*, a component of the SWI/SNF chromatin remodeling complex also known as *BICRA*, which increased in fall in 16 tissues (e.g. in cerebellar hemisphere; inset of Fig. 5A). Similarly, *RTF1*, a component of the RNA polymerase II transcription-associated *PAF1* complex, decreased in fall in nine tissues (Fig. S9A). This complex is deeply conserved across eukaryotes, and it has been described to be involved in the regulation of flowering time in plants (Ko et al. 2011; Dorcey et al. 2012) and to be required for induction of heat shock genes in animals (Tenney et al. 2006; Alpsoy and Dykhuizen 2018). PAF1c has been proposed to establish an antiviral state to prevent infection by incoming retroviruses: in case of infection by influenza A strain H3N2, PAF1c associates with viral NS1 protein, thereby regulating gene transcription (Liu et al. 2011). Other examples include *C4A*, decreasing in spring in 21 tissues (Fig. S9B), which localizes to the histocompatibility complex, and *KRT1*, which decreases in 24 tissues in winter (Fig. S9C), and encodes a keratin gene that is a downstream effector of the corticotropin hormone release pathway (Slenter et al. 2018). *Krt1* mutations in mouse have been associated with various phenotypic defects, ranging from abnormal circulating interleukin (Roth et al. 2012) to aberrant pigmentation of the epidermis (McGowan et al. 2006). Seasonal pigment variation is well known in mammals from the northern hemisphere (Zimova et al. 2018), which might suggest a conserved role of the corticotropin release pathway in pigment seasonal variation.

Finally, we focused on a set of 192 genes (1.1% of all expressed genes) that exhibited the strongest quantitative seasonal expression differences (at least a two fold-change in expression in one tissue-season pair). These were usually highly tissue specific, since only seven of these genes belonged to the set of 1,370 genes with recurrent seasonal patterns. These genes were enriched for functions related to epidermal differentiation and immunity (Fig. 5C), consistent with previous results in blood cells (Dopico et al. 2015). In line with the enrichment for multiple immune-related functions, these genes also exhibited significant overlap with gene sets that have been associated with COVID-19, including genes whose expression changes upon SARS-CoV-2 infection and genes predicted to be functionally related to *ACE2* (Fig. S12), although *ACE2* itself did not show a strong seasonal pattern. In particular, 24 out of the top 200 genes among the latter predictions (12.5%) were strongly seasonal, with 20 of them being up-regulated in the intestine specifically in the winter.

### Seasonal variation of gene expression of hormone genes

Hormones (Watts 2020; Tendler et al. 2021) have been described to broadly regulate the body’s seasonal physiology (Hanon et al. 2008; Dardente et al. 2014; Lomet et al. 2018). Thus, to explore whether genes that encode peptide hormones undergo particularly strong seasonal changes, we used a list of 62 genes with hormone-encoding capability based on Mirabeau *et al*. (Mirabeau et al. 2007) (Methods, Table S15). Overall, we found 39 (63%) hormone genes to be seasonal in at least one tissue (Fig. S10), a significant depletion respect to the whole genome (*P* = 7.011 × 10^-14^, two-sided proportion test). However, five hormone genes were included in at least one high-confidence seasonal gene set (Fig. 5D). Among these, *POMC*, a well-known photoperiodic hormone, was seasonal in 26 tissue-season pairs, mainly in summer. *POMC* expression has been shown to be dependent on longer-term photoperiod in the Siberian hamsters (Bao et al. 2019). Other seasonal hormones also have well known roles, mainly in the cardiovascular system and growth: *UCN*, a corticotropin for stress response and appetite regulation (Cullen et al. 2001; Vandael and Gounko 2019) and *ADM*, which is important for vasodilation (Geven et al. 2018). *GNRH1*, the gonadotropin-releasing hormone 1, which influences seasonal changes in other mammals (Hart et al. 1984), is seasonal in two artery tissues: the aorta and the tibial one. Mutations in *GNRH1* have been shown to be related to ischemic heart disease, which shows a seasonal pattern (Bhatia et al. 2017; Schooling and Ng 2019). Interestingly, leptin (*LEP*), a hormone involved in seasonal food-seeking behaviour, thermoregulation (Fischer et al. 2020), and obesity (Srivastava and Krishna 2007; Cahill et al. 2013), was altered only in winter in three tissues: adipose visceral, nerve, and blood (Fig. S10). Hormone genes were found robustly expressed not only in the tissues where they are seasonal but in several tissues. The tissue in which the largest number of seasonal hormone genes were robustly expressed (median TPM ≥ 5) was the hypothalamus (19 genes, Fig. S11). Pituitary was also among the tissues with a large number of seasonal hormones (11), together with testis (13) and frontal cortex (12).

### Seasonal changes in brain cytoarchitecture

Various studies have shown seasonal histological variation in different brain regions from several mammals, including anterior cingulate cortex in shrews (Lázaro et al. 2018), dendritic spines in amygdala in response to short days (i.e. in fall) in white-footed mice (Walton et al. 2012) and the volume of suprachiasmatic nucleus in humans (Hofman and Swaab 1992). To investigate potential seasonal changes in human brain’s cytoarchitecture, we used gene expression profiles of cell-type specific markers for a variety of brain cell types (McKenzie et al. 2018): astrocytes, neurons, oligodendrocytes, microglia, and endothelial cells (Table S16). We found that astrocyte markers significantly increased their expression in fall and decreased in summer (Fig. 6; see Methods for statistical analysis). In particular, we observed significant increased expression of astrocyte markers in the hypothalamus and frontal cortex in the fall, and a decrease in the cerebellum and frontal cortex in the summer. Oligodendrocyte markers, in contrast, tended to decrease expression in the fall, in particular in nucleus accumbens and anterior cingulate cortex (Fig. 6). In winter, all oligodendrocyte markers increased their expression in nucleus accumbens. Moreover, we observed a significant increase in the expression of neuronal markers in winter, particularly in the hypothalamus and spinal cord, and a global decrease in the fall (Fig. 6). These results are thus consistent with previous histological studies in humans and other mammals (Hofman and Swaab 1992; Walton et al. 2012; Lázaro et al. 2018) that showed that the relative volume or cytoarchitecture of astrocytes, oligodendrocytes and neurons change with the season in a subregion-specific manner.

**Fig. 6:**
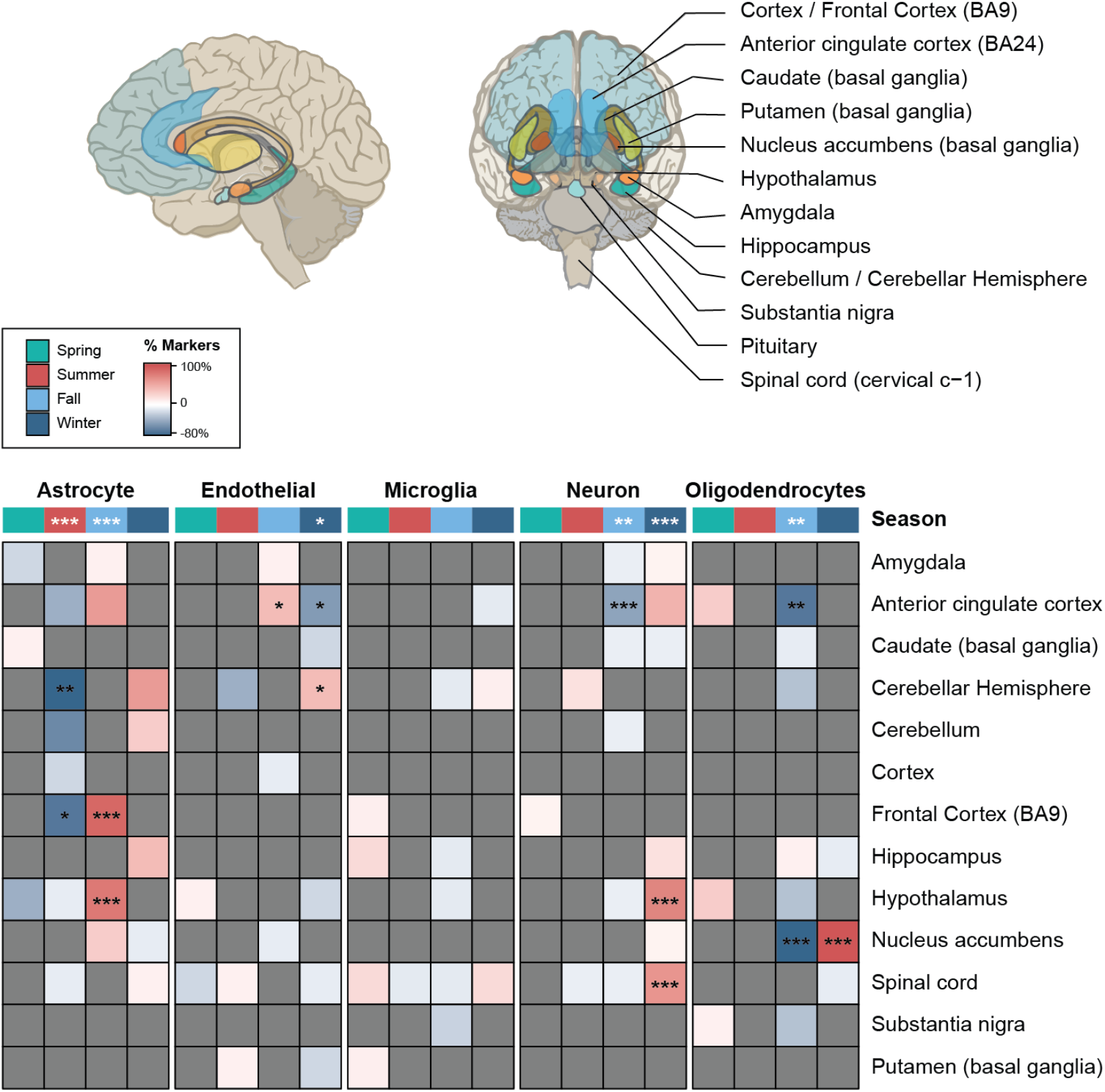
Seasonal variation in cell-type specific markers across human brain regions. Percentage of the marker genes down-regulated (blue) or up-regulated (red) for five brain cell types: astrocyte, endothelial, microglia, neuron, and oligodendrocytes, in brain subregions (depicted in the brain scheme above, downloaded and modified from the GTEx web portal). All significant markers for a cell type for a specific season in a subregion were differentially expressed in the same direction, over or under expressed. Markers without significant effects were colored in grey. P-values were obtained by randomizing the observed number of up- and down-regulated markers per cell type across regions and seasons conservatively allowing only up- or down-regulated makers for a given region-season combination (see Methods for details). *** *P* < 0.001, ** 0.001 ≤ *P* < 0.01, ^*^ 0.01 ≤ *P* < 0.05.

## Discussion

By leveraging the rich transcriptome data produced by the GTEx project, we have investigated the impact of the circadian and circannual cycles in the human transcriptome across multiple tissues. Ruben *et al*. (Ruben et al. 2018) has also recently used the GTEx data to investigate circadian variation in gene expression; however, in that case, the time of death was inferred from the expression of known circadian genes. Here, in contrast, we use the actual time of death reported in the GTEx metadata. Even though GTEx captures a single snapshot of the tissue transcriptomes of the donors at the time of death, since this time is approximately uniformly distributed throughout the day and the year, the aggregation of the data snapshots across individuals produces trajectories that allowed us to investigate temporal variations in gene expression. Moreover, although there is an impact of the death of the individual in the transcriptome, which is tissue-specific, this can be properly controlled for (Ferreira et al. 2018). The GTEx medatada, however, by making accessible only the time of the day and the season of death of the donors makes this investigation challenging. We have addressed this limitation by discretizing the continuous circadian variation into day versus night, and the circannual variation into seasons. This impacts the way in which we specifically formulate the questions, i.e. we do not refer to the variations reported here as circadian and circannual, but as day-night and seasonal, respectively, and restricts the statistical methods that can be employed to analyze the data. Thus, while ANOVA would appear the natural approach to identify seasonal changes in gene expression, the large variation in the date of death within each season decreases in practice the power to detect significant changes in gene expression; therefore, we opted for focusing on a method that allows leveraging the unknown date of death and that could be more directly compared with that employed for the day-night analyses (i.e. performing pairwise differential gene expression analyses). Despite these and other caveats (e.g. effect of artificial lights, ethnic origins, date of the RNA extractions, etc.), our approach was able to properly capture at least part of the real circadian and circannual transcriptional variation, since we have been able to recapitulate previous findings regarding day-night variation.

We performed an initial comparison of both types of variation within and among tissues using relaxed cut-offs, finding that the effect of day-night variation in gene expression was comparable to that of the seasonal cycle, but affecting different genes and tissues. Remarkably, day-night variation in gene expression was more prominent in liver, lung, heart, and upper digestive tract, reflecting the involvement of the organs of the thoracic cavity in circadian processes (Nosal et al. 2020), while seasonal variation had the strongest effect in brain subregions and testis, mirroring the role of the brain-gonadal hormonal axis in regulating the physiological responses to seasonal variation (Kang et al. 2020). Moreover, we showed that the effect of day-night and/or seasonal variation for most genes was highly tissue-specific. In the case of the day-night genes, this is in agreement with the hypothesis that one molecular clock program is present in all tissues, but the circadian processes triggered downstream are highly tissue-specific (Scheiermann et al. 2013). However, subsets of genes showed highly consistent up- or down-regulation during day or night or in specific seasons across multiple tissues, which we defined as high-confidence day-night or seasonal genes, respectively.

Importantly, the day-night high-confidence set was highly enriched for genes with a known circadian pattern and included most known core clock genes. In addition, various metabolic functions were enriched among day or night genes, in line with multiple reports (e.g. (Neufeld-Cohen et al. 2016; Hatori et al. 2012)). The direction of the change in gene expression for the core clock genes and other high-confidence day-night genes was highly consistent among tissues, but we found two cases in which changes in gene expression occurred unexpectedly in opposite directions between brain and non-brain tissues. This is the case of *NR1D2* and *RGS1*, in which the day-night changes in gene expression occurred in the opposite direction in the brain than in tissues from the rest of the body. This suggests that the cellular environment could modulate the interpretation of the core clock signals, and produce different (or even opposite) outputs in different tissues in response to the same environmental cues.

Among all core clock genes, *ARNTL* and *NPAS2* had a differential day-night transcriptional behaviour across the largest number of tissues. In the circadian negative feed-back loop, *ARNTL* forms an heterodimer with either *NPAS2* or its paralog *CLOCK*, positively regulating the circadian pattern (Scheiermann et al. 2013). Interestingly, we found that *NPAS2* shows day-night gene expression differences in a much larger number of tissues than *CLOCK* (30 vs. 12 tissues), suggesting a more relevant role for *NPAS2* in the circadian modulation across tissues in human. Similarly, *NPAS2* was detected as cycling in 23 baboon tissues compared to eight tissues for *CLOCK* (Mure et al. 2018). In contrast, mouse *Clock* and *Npas2* cycled in a similar number of tissues (eight and seven out of 12 tested tissues, respectively) (Li et al. 2018). Whether this difference is a lineage-specific divergence between primates and rodents, or it may be related to distinct diurnal-nocturnal habits (diurnal for human and baboon, nocturnal for mouse) will require investigation in other mammalian groups. In the latter scenario, however, the differential use of *NPAS2* and *CLOCK* could contribute to establishing the opposite circadian patterns across tissues that we observed between primates and mouse.

At the seasonal level, our results highlight the importance of the brain-gonadal hormonal axis. Many physiological and behavioral changes across seasons such as breeding, mating, molting, foraging, and hibernation (Walton et al. 2011; Dardente et al. 2014; Ebling 2014; Watts 2020) are known to be regulated by the endocrine system in mammals and birds, especially by the brain-thyroid and brain-gonadal axes, and can elicit different reactions depending on the receiving tissues. This is reflected in the tissue-specific seasonal variation in gene expression that we have uncovered, which affects prominently testis and many brain regions. When investigating specifically the expression patterns of genes involved in functions related to the hormonal axis, we found these to be predominantly seasonal in core tissues from the endocrine system, such as hypothalamus and pituitary. Surprisingly, however, we found that hormone genes are overall depleted for seasonal variation.

Infectious diseases are well-known to have a seasonal pattern, as illustrated, for instance, by the current COVID-19 pandemics or the yearly peaks of flu infections. Consistently, we have observed a significant enrichment for immune related functions among both the winter high-confidence gene set (Fig. 5B) as well as the strongly seasonal genes (Fig. 5C). Moreover, we have found a significant number of genes in various COVID-19 related pathways and associated gene sets to be strongly seasonal in a tissue specific manner. In particular, these included genes that change expression in response to SARS-CoV-2 infection and genes predicted to be functionally related to *ACE2*, an endogenous membrane protein that mediates SARS-CoV-2 infection. These genes are likely not specific to SARS-CoV-2 infection, and their strong seasonality revealed here could shed light on the molecular mechanisms underlying the seasonality of viral infections in general.

Finally, by evaluating cell type-specific markers, our results suggest a substantial seasonal remodeling of the cytoarchitecture of certain human brain areas. We found a general increase of astrocytes in fall and winter and a decrease in summer across many subregions. These changes in the expression of cell type markers, similar to the volumetric changes described in other mammals, were subregion-specific. Interestingly, we found a reduction of neuronal markers in the anterior cingulate cortex in fall, a subregion whose neuronal soma and dendrite size shrank in the cold season in shrews (Lázaro et al. 2018). Whether or not these and other putative cytoarchitectural changes are conserved in other mammals, which are their functional implications, and how they might contribute to the seasonal patterns of some psychiatric and brain diseases (Philpot et al. 1989; Joseph-Vanderpool et al. 1991; Owens and McGorry 2003; Walton et al. 2012; Lim et al. 2017) need to be further investigated.

In summary, our work expands our understanding of the transcriptional impact of the physiological changes associated with the day-night cycle and, for the first time across multiple human tissues, of the seasonal cycle. Moreover, these results constitute a large resource for the community to further investigate the impact of day-night and seasonal variation in the human transcriptome.

## Methods

### RNA-seq datasets

The RNA-seq data was generated, mapped and quantified by the GTEx consortium (GTEx v8) (GTEx Consortium 2020). Tissues with less than 100 donors were discarded from the analyses (Kidney - Medulla, Kidney - Cortex, Cervix - Ectocervix, Fallopian Tube, Cervix - Endocervix, and Bladder) as well as two cell lines (Cells - EBV-transformed lymphocytes and Cells - Cultured fibroblasts). For the Whole Blood tissue, all pre-mortem samples were discarded from the analysis for homogeneity with the other tissues. After filtering for samples with available covariates for the differential analyses, we employed 16,151 samples from up to 46 tissues for 932 individuals. GTEx individuals are biased toward old males (median age = 55 years, 67% male). GENCODE v26 (Harrow et al. 2012) was used for GTEx as well as the annotation for the protein-coding genes.

### MetaCycle

The function *meta2d* from the R package MetaCycle (v.1.2) was runned for each 46 tissues separately using the default parameters. Genes with a median TPM < 1 per tissue were filtered out and the time of death of the individuals were converted into hours.

### Classification of the time of death as day or night and by season

Circadian patterns had been previously analyzed using the GTEx data, but inferring the time of death from the expression of marker genes (Ruben et al. 2018; Anafi et al. 2017). Here we use, instead, the actual time of death as provided by the GTEx consortium. Using this time of death, individuals had been classified between dead during the day or dead during the night if their time of death was falling into the following intervals [08:00-17:00) and [21:00-05:00) respectively (Fig. 1A). Other times of death have been discarded for the day-night analysis to avoid taking into account any RNA-seq samples coming from people where the day status was unsure, i.e. twilight. Using this classification, 351, 315 and 266 individuals have been classified as day, night and twilight respectively, of which only the 666 day and night individuals (11,527 samples) were used for differential expression for the day-night cycle. The time corresponding to day and night was manually curated from the Boston (Massachusetts) sunrise and sunset intervals during the year (using https://www.timeanddate.com/sun/usa/boston). The season of death was provided by the GTEx consortium with 190, 221, 282, and 239 donors that died in spring, summer, fall, and winter respectively. Note that for the season, no people were discarded regarding the time of death. We also provide the information about the month in which the RNA was extracted for each season of death (Table S17).

### Differential gene expression between day and night

Differential expression between day and night was performed separately on the 46 tissues using samples from the 666 individuals classified as day or night, ranging from 98 samples (Uterus) to 560 samples (Muscle - Skeletal). Genes were filtered per tissue, removing all genes with a median TPM < 1 over the day and night samples, leading to 18,022 protein-coding genes expressed in at least one tissue (31,530 genes including all biotypes). The analyses were run using R v3.6.1 (Team 2019), the TMM normalisation method from *edgeR* (with the *calcNormFactors* function) (Robinson et al. 2010; McCarthy et al. 2012), and the *voom-limma* pipeline (with the *voom, lmFit*, and *eBayes* functions) (Law et al. 2014; Ritchie et al. 2015) using default parameters. The significance of the time of death was assessed correcting for the following covariates: sex if the tissue was not sex-specific; age; body mass index (BMI); the postmortem interval; the season of death. All genes with an associate *P* ≤ 0.05 and an absolute log_2_ fold-change ≥ 0.1 were considered as day-night. Results, including all biotypes, are available in Supplementary Dataset 1 (including p-values, adjusted p-values and log_2_ fold-changes). The number of differentially expressed genes across experiments was not affected by the number of samples (Fig. S13A) and no effect was observed regarding the ratio of day/night samples and the number of differentially expressed genes between day and night (Fig. S13B). The volcano plots of the analyses for each tissue are available in Supplementary Dataset 3.

### Season-specific gene expression

Differential expression between seasons was performed separately on the 46 tissues using samples from the 932 individuals, ranging from 139 samples (Brain - Substantia nigra) to 789 samples (Muscle - Skeletal). Genes were filtered as described in the section ‘Differential expression between day and night’. This resulted in a set of 18,018 protein-coding genes expressed in at least one tissue (31,517 genes including all biotypes). The effect of each season was assessed by comparing one season against all the others, leading to four differential expression analyses, one for each season (see ‘Differential expression between day and night’ for details). This approach was taken to make the analyses more consistent with that of day-night variation (i.e. using the exact same method), and to minimize the impact of the fact that only the season, but not the actual day, of death is known in GTEx (i.e. two individuals could have died one day apart but in two different seasons, or 90 days apart but still within the same season). The covariates used were the following: time of death classified as: day, night or twilight; sex if the tissue was not sex-specific; age; BMI; postmortem interval. All genes with an associate *P* ≤ 0.05 and an absolute log_2_ fold-change ≥ 0.1 were considered as seasonal. Results, including all biotypes, are available in Supplementary Dataset 2 (including p-values, adjusted p-values and log_2_ fold-changes). The number of seasonal genes was not related to the number of samples per the tissue (Fig. S14A) and no relation was observed between the proportion of samples available for a given season over the total and the ratio of up- and down-regulated genes. (Fig. S14B). The volcano plots of the analyses for each tissue-season pair are available in Supplementary Dataset 3.

### Testing tissues for day or night

To test if day-night genes in two tissues or more are more prone to be either day or night, we computed for each gene the ratio of day tissues over the total number of tissues for this gene. Genes will be considered to have a “consistent” ratio if their ratio is < 0.25, mainly night tissues, or > 0.75, mainly day tissues. For each number of tissues, from two to ten tissues, we calculate the percentage of genes that are considered to have a consistent ratio and the expected probability to be in the consistent ratio distribution following a binomial distribution with a probability of success (day tissue) equal to 0.5 and a number of draws equal to the number of genes that are day-night for this number of tissues. We then performed a binomial test for each number of tissues. The results are available at Table S4.

### Definition of the high-confidence gene sets

To define the five high-confidence gene sets (one for day-night genes and four for the seasonal genes), we perform three binomial tests per gene: i) using all tissues, ii) using only non-brain tissues, and iii) using only brain regions. The number of draws was the number of tissues in which the gene was day-night or seasonal and the number of successes the number of tissues in which the gene was day or up. Only two of the tested genes have divergent behavior between non-brain and brain tissues: *NR1D2* (11 night non-brain tissues and 9 day brain tissues) and *RGS1* (6 day non-brain tissues and 6 night brain tissues). No divergent behavior was observed for the seasonal genes.

### Comparison of human, baboon and mouse core clock genes

The baboon results were downloaded from Mure *et al*. (Mure et al. 2018). We extracted the significant genes using the same threshold as Mure *et al*. (*P* ≤ 0.05). Common tissues between GTEx and baboon have been manually curated (Table S6). The mouse circadian genes were downloaded from the CirGRDB database (Li et al. 2018), which include genes already defined as circadian, and we only selected the genes found by RNA-seq and from the publication with the PubMed ID 25349387, corresponding to the publication by Zhang *et al*. (Zhang et al. 2014). Common tissues between GTEx and the mouse tissues list had been manually curated (Table S7). This dataset was used to compute the number of occurrences of *Clock* and *Npas2* in mouse tissues.

To perform the comparison of the core clock genes in human vs baboon, we extracted the core clock genes in both species in the 20 common tissues between the GTEx and the baboons’ tissue. From this, we analyzed only genes that were significant in both species, i.e. we discarded the genes found significant in either baboon or human but not in both. The same analysis was done to compare human and mouse.

### Functional enrichment analyses

The high-confidence sets for day-night and seasonal genes, separated by day/up- and night/down-regulated genes, were used as input for the enrichment analysis tool Enrichr (Chen et al. 2013; Kuleshov et al. 2016). Results for GO Biological Process were retrieved and are provided fully in Tables S9 (day-night) and S14 (seasons). To simplify visualizations in Fig. 3D and 5B,C, we disregarded GO terms with fewer than 10 total genes. In addition, we removed redundant categories by excluding those terms sharing at least 80% of genes with a category with a lower p-value. For COVID-19 related gene sets, gene sets with an adjusted p-value greater than 0.05 were discarded.

### Circadian and hormone gene sets

The protein sequences, along with the IDs, of proteins annotated as circadian, either experimentally or by orthology, in human were downloaded from the Circadian Gene DataBase website (http://cgdb.biocuckoo.org/) (Li et al. 2017b) the 01/26/2021. The correspondence between the Ensembl protein ID and the UniProtKB ID with the Ensembl gene ID was downloaded using BioMart from Ensembl (Smedley et al. 2015). The list of hormone genes was downloaded from Mirabeau *et al*. (Mirabeau et al. 2007). The Ensembl peptide IDs were linked to their respective Ensembl gene ID using R and the *biomaRt* package (Durinck et al. 2005, 2009). The gene IDs for hormones with deprecated peptide IDs were retrieved manually using the Ensembl website (http://www.ensembl.org) and the ones that were obscelets were removed. Two hormone genes were added manually: *GH1* and *LEP*. The list is available at Table S15.

### Statistical assessment of cell-type specific marker changes across seasons

For each cell type, we evaluated a fixed number of markers (Astrocyte: 10, Endothelial: 10, Microglia: 9, Neuron: 11, Oligodendrocytes: 10) from (McKenzie et al. 2018) (Table S16). We assessed the significance of the number of up- or down-regulated markers for each cell type in each specific region-season pair (each cell in Fig. 6) as well as for all regions together (top row in Fig. 6). For this purpose, we first counted the total number of instances in which markers for a given cell type are up and down-regulated across the entire set of region-season pairs (Astrocyte: 42 up, 36 down; Endothelial: 12 up, 20 down, Microglia, 9 up, 8 down; Neuron, 27 up, 12 down; Oligodendrocytes, 18 up, 27 down). Then, we performed 1,000,000 randomizations of the up and down instances across seasons and regions. We conservatively allowed only up or down markers in a given region-season pair for each cell type, since this is what we have observed in the real data (i.e. no cell type markers showed contradictory patterns in any region-season pair). To calculate the p-value for each region-season pair we simply obtained the number of randomizations in which the number of up- or down-regulated markers was equal or higher than the tested region-season pair and divided it by one million. These p-values were Bonferroni corrected for multiple testing (260 tests). To calculate the p-value across regions, we sum all up and down instances across regions, and again obtained the number of randomizations in which the number of up- or down-regulated markers was equal or higher than the sum across regions and divided it by one million. These p-values were also Bonferroni corrected for multiple testing (20 tests).

### Data access

All GTEx open-access data are available on the GTEx Portal (https://gtexportal.org/home/datasets). All GTEx protected data are available via dbGaP (accession phs000424.v8). Access to the raw sequence data is now provided through the AnVIL platform (https://gtexportal.org/home/protectedDataAccess).

## Acknowledgments

We thank the donors and their families for their generous gifts of biospecimens to the GTEx research project; the Genomics Platform at the Broad Institute for data generation. The Genotype-Tissue Expression (GTEx) project was supported by the Common Fund of the Office of the Director of the National Institutes of Health (https://commonfund.nih.gov/GTEx). Manuel Muñoz-Aguirre and Thomas Derrien for their feedback on the manuscript. Diego Garrido Martín for his help on side analysis. Claudia Vivori and Vanessa Vega Méndez for providing help with graphical presentations.

## Authors’ contributions

V.W. Design and creation of the study, major data analysis and visualization, final main and supplementary data and figure production and management, writing of the methods, and revisions. R.S. Design and creation of the study, writing of the main text, major data analysis, questions, interpretations, and visualization of earlier editions. R.A. data visualization and analysis, and helpful discussions. M.I. revisions, intellectual contribution, and final figure production. R.G. revisions, accommodation of the research, and intellectual contribution.

