## Supplementary Figures for "Day-night and seasonal variation of human gene expression across tissues"

**Supplementary Table Legends**

**Table S1 - Day-night and MetaCycle results.** For each tissue, number of genes found by the day-night analysis, MetaCycle, and the genes they found in common.

**Table S2 - Phase classification of the genes missed by the day-night analysis but found by MetaCycle.** For each tissue, for the genes that are found by MetaCycle but missed by the day-night analysis, the number of genes that are classified as day, night, twilight, or within one hour of the twilight based on the predicted phase of MetaCycle.

**Table S3 - Summary statistics of day-night expression analysis**. For each tissue, number of genes with differentially upregulated expression during the day or during the night, percentage of expressed genes with a day or night expression, number of expressed genes, and log2 of the ratio day vs night upregulation. Corresponding to Fig. S2A.

**Table S4 - Binomial test of the number of day-night genes in consistent bins.** For genes that were identified as day-night in a given number of tissues (from two to ten), it is shown: the number of genes that are in a consistent bins, i.e. genes with a ratio of day tissues over the total number of tissues $< 0.25$ or $>0.75$, the total number of genes, and the p-value from the binomial test obtained using the expected ratio from a random distribution for that number of tissues as the hypothesized probability of success ("Probability of being in consistent bins"). See Methods for details.

**Table S5 - List of core clock genes**.

**Table S6 - Homologous tissues for human and baboon**. Compared homologous tissues between the human GTEx dataset and baboon dataset from Mure *et al.*

**Table S7 - Homologous tissues for human and mouse**. Compared homologous tissues between the human GTEx dataset and mouse dataset from Li *et al.*

**Table S8 - Day-night genes and their associated binomial p-value.** For each gene, the number of tissues for which the gene is day or night, for i) all tissues, ii) non-brain tissues, and iii) brain tissues. For each of the previous tissue classification, a binomial p-value is associated to test if the gene has a statistically significant preference to be highly expressed during the day or the night. Genes with a $P \leq 0.05$ for at least one of the tissue classifications are part of the high-confidence day-night gene sets. Note that for a gene to reach a $P \leq 0.05$ it needs to have been identified day-night in at least 6 tissues. See Methods for details.

**Table S9 - Gene Ontology enrichment results for the high-confidence day-night genes for day and night genes separately.** Enrichment performed with EnrichR. See Methods for details.

**Table S10 - Genes associated with sleep traits**. List of 254 protein-coding genes expressed in GTEx and that were previously reported to increase the risk of insomnia or to be associated with other sleep traits in humans.

**Table S11 - Summary statistics of the seasonal expression analysis**. For each tissue and season, number of genes with differentially up- or down-regulated expression, percentage of expressed genes with up or down regulation in that given tissue and season, number of expressed genes, and log2 of the ratio up/down.

**Table S12 - Overlap between day-night and seasonal variation**. For each tissue, number of genes with day-night expression, seasonal expression, overlap between the two sets and p-value for the overlap (Fisher's Exact test with and without Benjamini - Hochberg multiple testing correction).

**Table S13 - Seasonal genes and their associated binomial p-value.** For each season and seasonal gene, the number of tissues for which the gene is up- or down-regulated, for i) all tissues, ii) non-brain tissues, and iii) brain tissues. For each of the previous tissue classification, a binomial p-value is associated to test if the gene has a statistically significant preference to be up- or down-regulated in that season. Genes with a $P \leq 0.05$ for one of the tissue classifications are part of the high-confidence seasonal gene sets. Note that for a gene to reach a $P \leq 0.05$ it needs to have been identified as differentially expressed in that season in at least 6 tissues. See Methods for details.

**Table S14 - Results from the Gene Ontology enrichment analysis for the high-confidence seasonal genes per season and for up- and down-regulated genes separately.** Enrichment performed with EnrichR. See Methods for details.

**Table S15 - List of human hormone genes.** List of 62 genes with hormone-encoding capability based on Mirabeau *et al.*

**Table S16 - Cell type specific markers**. List of genes serving as cell type specific markers in the human brain from McKenzie *et al.*

**Table S17 - Month of RNA extraction and season of death.** The month of the RNA extractions of the samples and the season of death of the individuals.

**Supplementary Dataset 1 - Raw data for day-night gene expression variation**. For each human gene, results of the differential gene expression analysis between day and night. These data can be downloaded using the following public link: https://www.dropbox.com/s/7vbytq26fd6jjeg/Supplementary_Dataset_1.xlsx?dl=0

**Supplementary Dataset 2 - Raw data for seasonal gene expression variation**. For each human gene, results of the differential gene expression analysis for each season against the rest. These data can be downloaded using the following public link: https://www.dropbox.com/s/1leckfcmry21ly2/Supplementary_Dataset_2.txt.gz?dl=0

**Supplementary Dataset 3 - Volcano plots for differential gene expression in day-night and seasonal variation**. For each tissue, volcano plot showing −log10(p−value) vs log2(fold-change) for day-night as well as for each season comparison. These plots can be downloaded using the following public link: https://www.dropbox.com/s/8yubqw0ezcqcvi3/Supplementary_Dataset_3.zip?dl=0

**Figures**

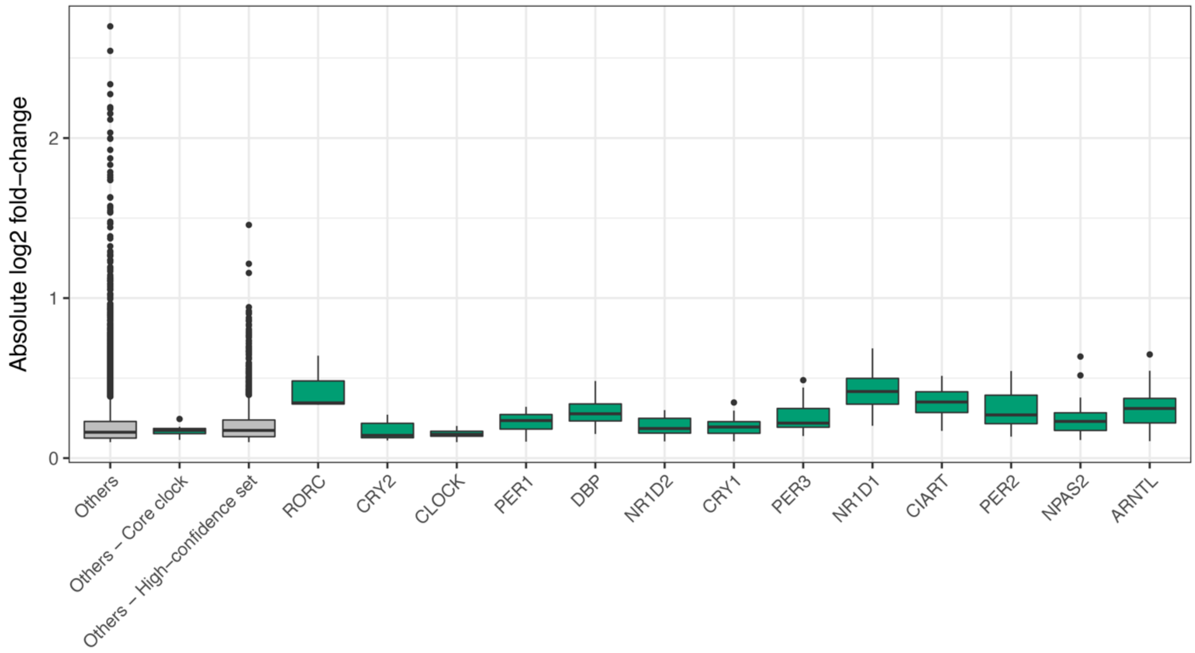


**Fig. S1 - Absolute *log₂* fold-change (or effect size) of day-night genes.** Each core clock gene in the high-confidence day-night set is plotted separately (green boxplots). The rest of the genes are pooled together in 'Others - Core clock' if they are in the list of the core clock genes, in 'Others - High-confidence set' if they are part of the high-confidence set, or in 'Others' otherwise.

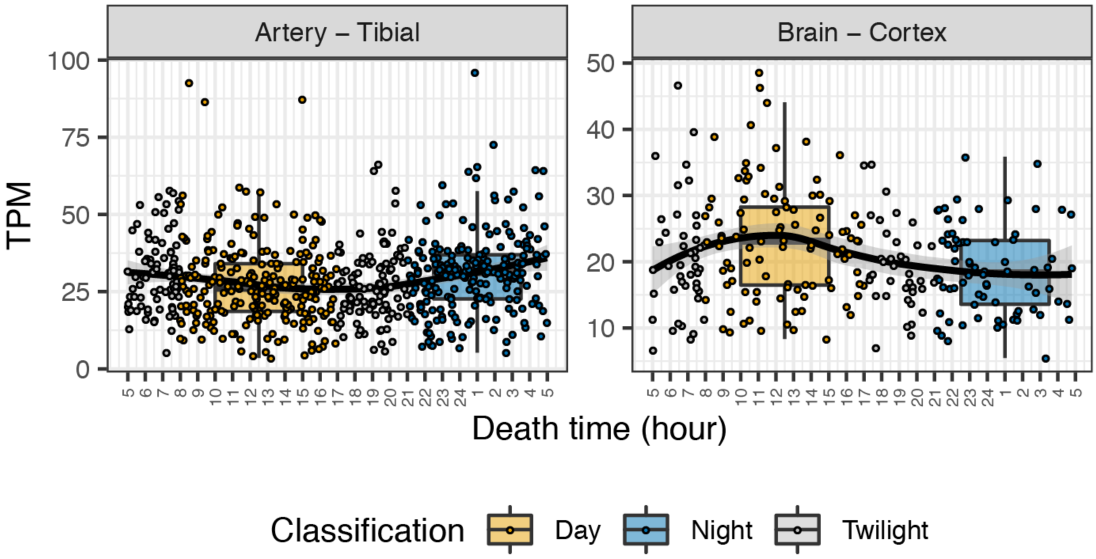


**Fig. S2 - Contrasting *NR1D2* circadian behaviors in brain and other tissues.** TPM values for *NR1D2* in the artery - tibial (left) and in the brain - cortex (right). The colors of the dots represent the classification of the individuals according to the time of death of the donor: during the day (yellow), during the night (blue), or in-between for twilight (grey). The samples classified as twilight have been discarded for the day-night analysis. The "circadian" curve was created using the *geom_smooth* function from *ggplot2* in R with the 'loess' method.

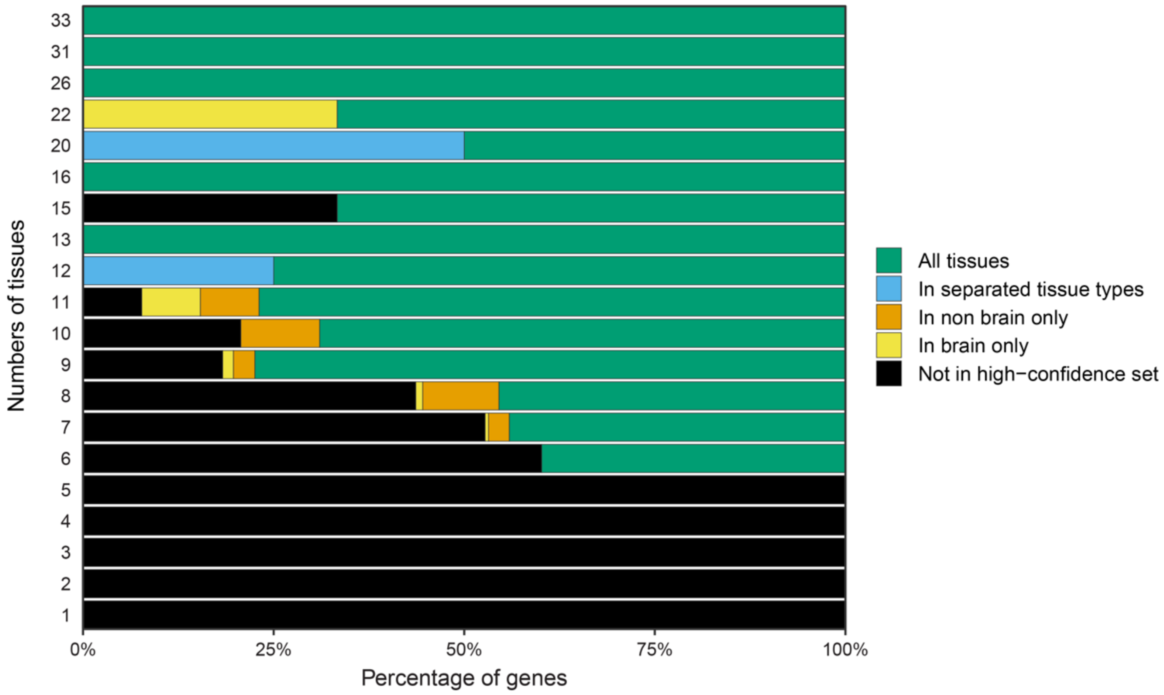


**Fig. S3 - Classification of genes in the high-confidence day-night set.** For each number of tissues in which a gene was identified as day-night, the percentage of genes that are consistently up-regulated either during day or during night for: i) all tissues taken together ($P \leq 0.05$ for the binomial test on all tissues; All tissues, green), ii) in non-brain and in brain tissues separately ($P \leq 0.05$ for the binomial test on non-brain and brain tissues and $P > 0.05$ for all tissues; In separated tissue types, blue), iii) in non-brain tissues only ($P \leq 0.05$ only for the non-brain tissues test; In non-brain only, orange), iv) in brain tissues only ($P \leq 0.05$ only for the brain tissues test; In brain only, yellow), and v) inconsistent between day and night tissues ($P > 0.05$ for all binomial tests; Not in high-confidence set, black). See Methods for details.

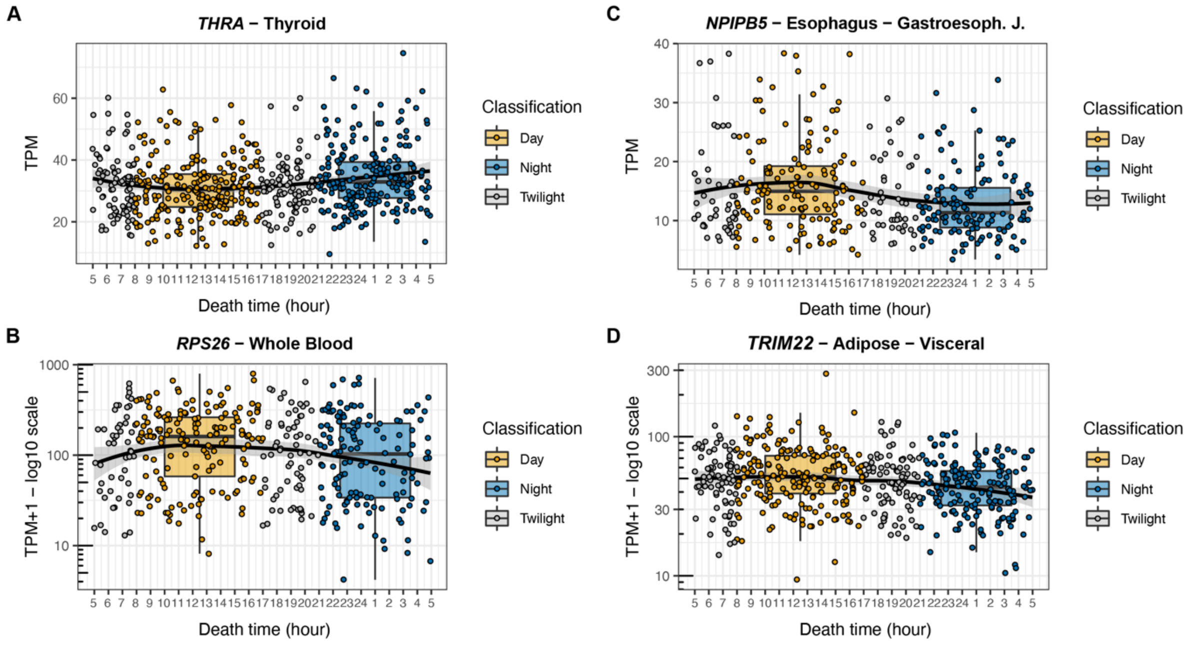


**Fig. S4 - Examples of day-night genes.** Expression values (TPM) values for three high-confident day-night genes at the time of death of the GTEx donors: (A) *THRA* in thyroid, (B) *RPS26* in the whole blood, (C) *NPIPB5* in the esophagus - gastroesophageal function, and (D) *TRIM22* in the adipose - visceral (omentum). The colors of the dots represent the classification of the individuals according to their time of death of the donor: during the day (yellow), during the night (blue), or in-between for twilight (grey). The samples classified as twilight have been discarded for the day-night analysis. The "circadian" curve was created using the *geom_smooth* function from *ggplot2* in R with the 'loess' method.

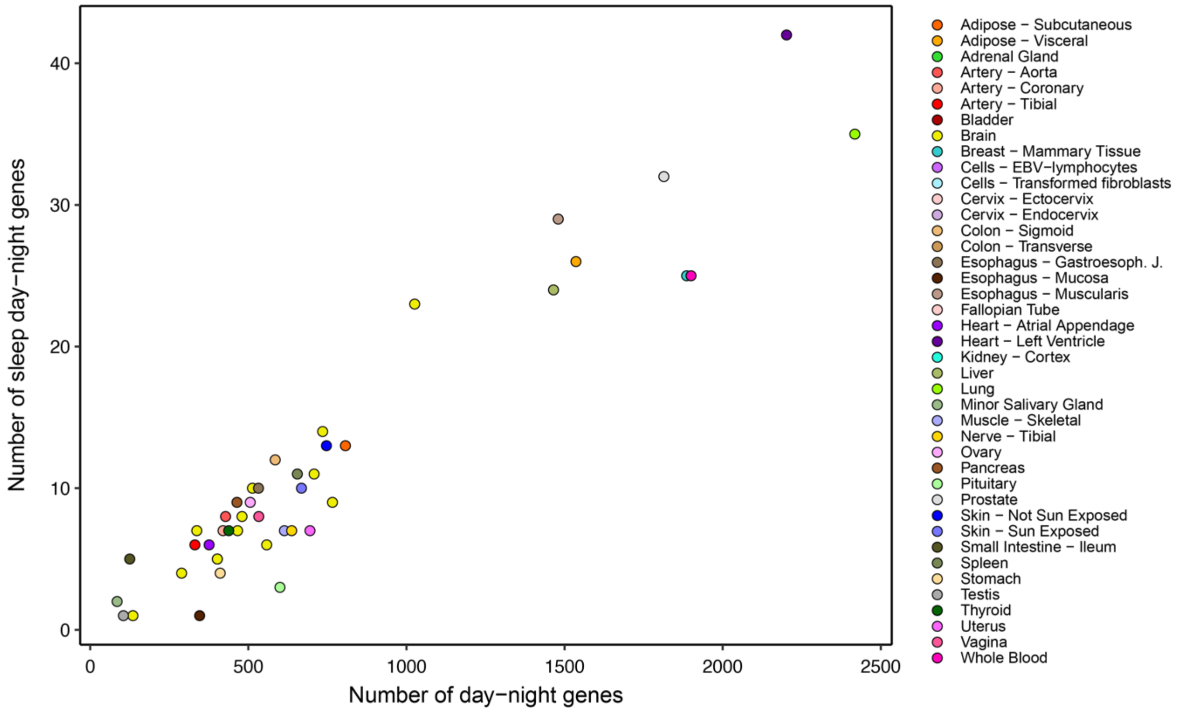


**Fig. S5 - Day-night variation of sleep associated genes.** Number of sleep day-night genes (y-axis) vs. total day-night genes (x-axis) per tissue.

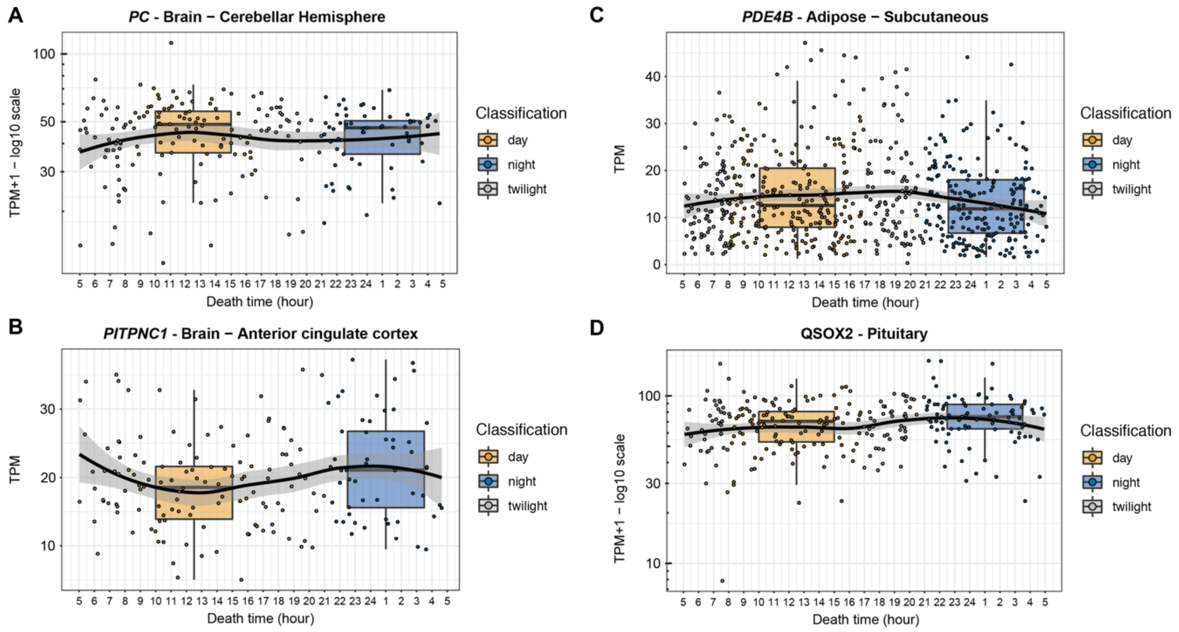


**Fig. S6 - Example of day-night sleep genes.** Expression values (TPMs) for the four sleep-related genes in the day-night high-confidence gene set and annotated in the Circadian Gene DataBase: (A) *PC* in cerebellar hemisphere, (B) *PITPNC1* in anterior cingulate cortex, (C) *PDE4B* in subcutaneous adipose, and (D) *QSOX2* in pituitary at the time of death of the GTEx donors. The colors of the dots represent the classification of the individuals according to the time of death of the donor: during the day (yellow), during the night (blue), or in-between for twilight (grey). The samples classified as twilight have been discarded for the day-night analysis. The "circadian" curve was created using the *geom_smooth* function from *ggplot2* in R with the 'loess' method.

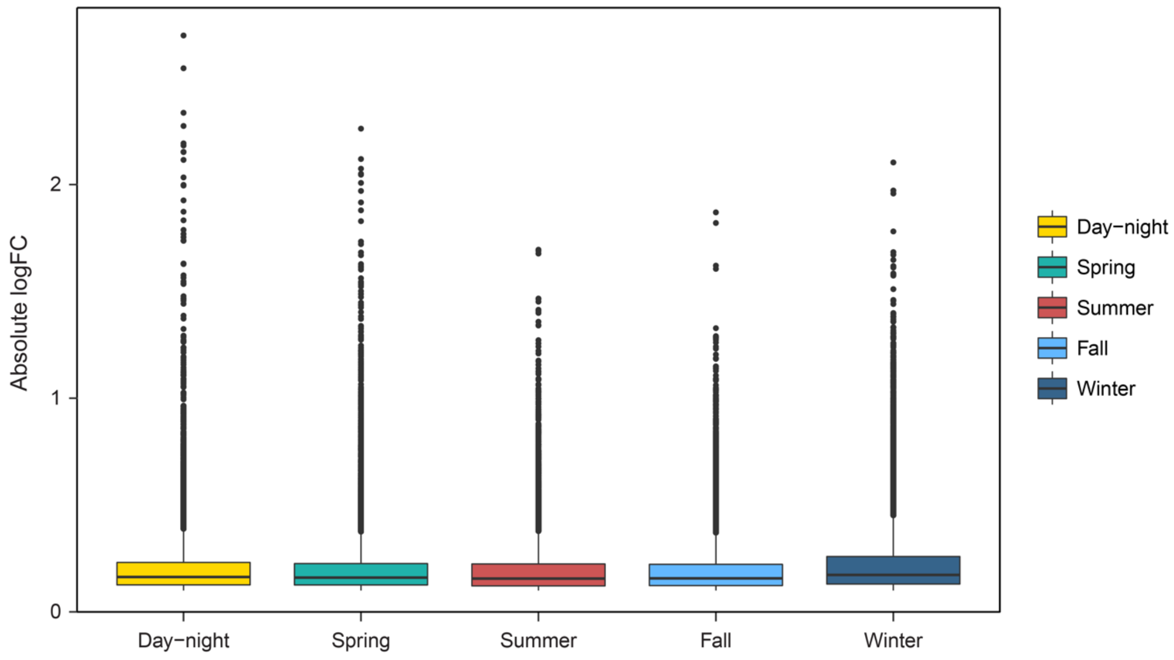


**Fig. S7 - Effect size of day-night and seasonal variation in gene expression.** Boxplots showing the absolute $log_2$ fold-change (or effect size) of day-night genes (yellow) or seasonal genes (red: summer, green: spring, light blue: fall, dark blue: winter).

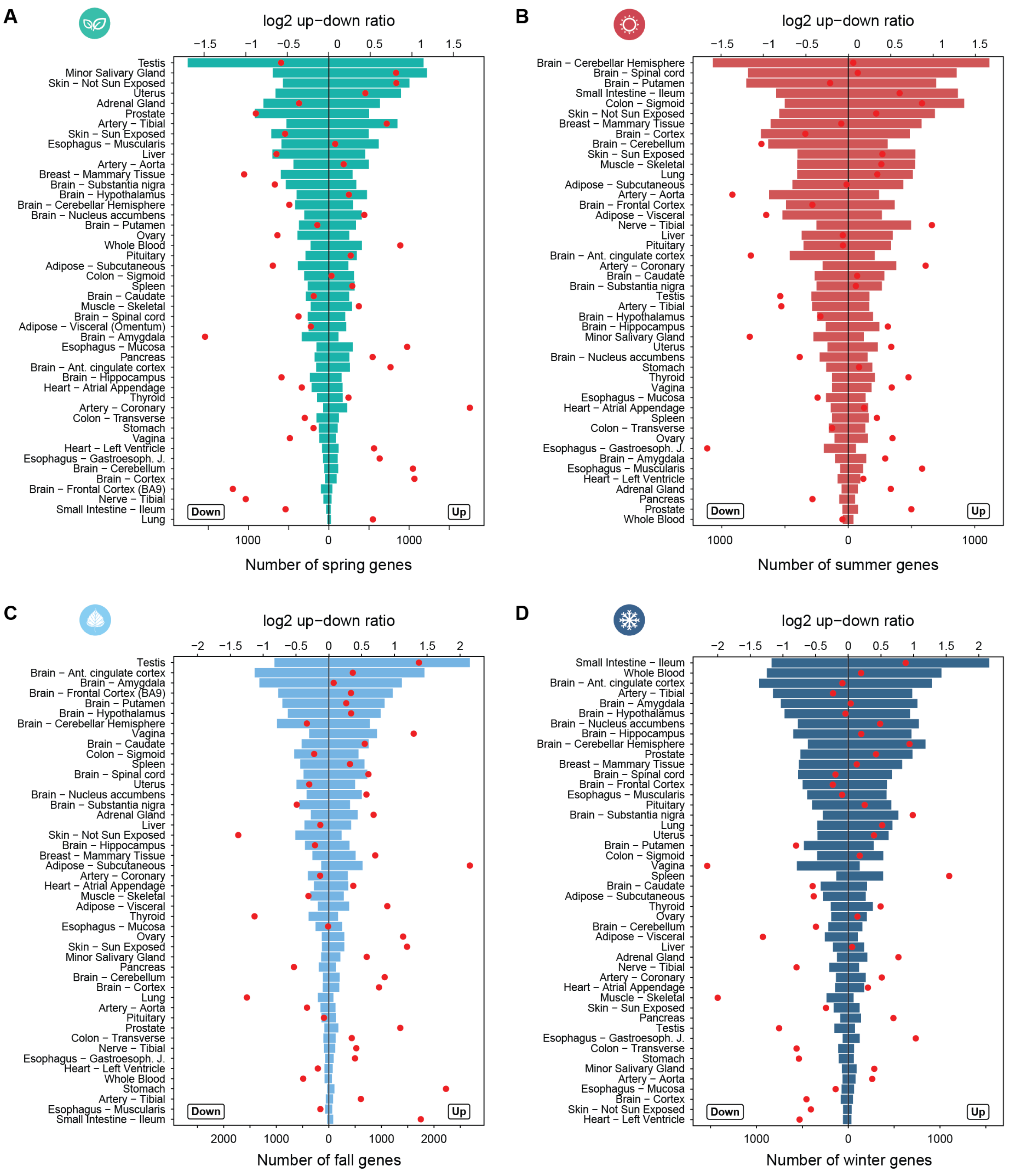


**Fig. S8 - Landscape of seasonal gene expression variation by tissue.** Number of over-expressed (Season up, right side) and under-expressed (Season down, left side) seasonal genes per tissue in GTEx (bottom axis) for each season, spring (A), summer (B), fall (C), and winter (D). The tissues were ordered according to the total number of seasonal genes per season. The red dot represents the $\log_2$ ratio between the number of over and under genes (top axis).

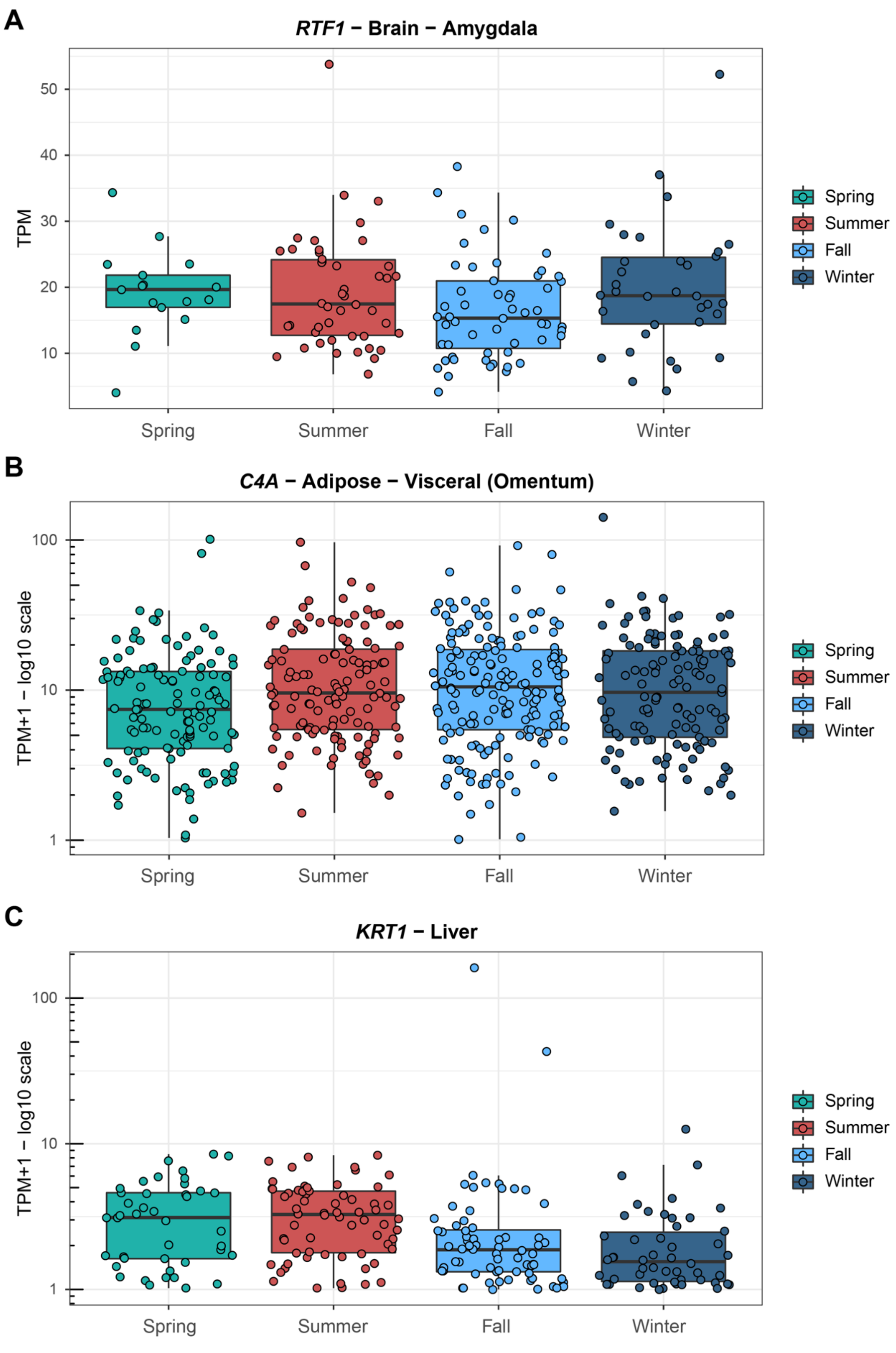


**Fig. S9 - Examples of seasonal genes.** Boxplot of the TPM values on a $\log_{10}$ scale of three top seasonal genes in a tissue where they are differentially expressed: *RTF1*, under-expressed in fall (A), *C4A* under-expressed in spring (B), and *KRT1* under-expressed in winter (C).

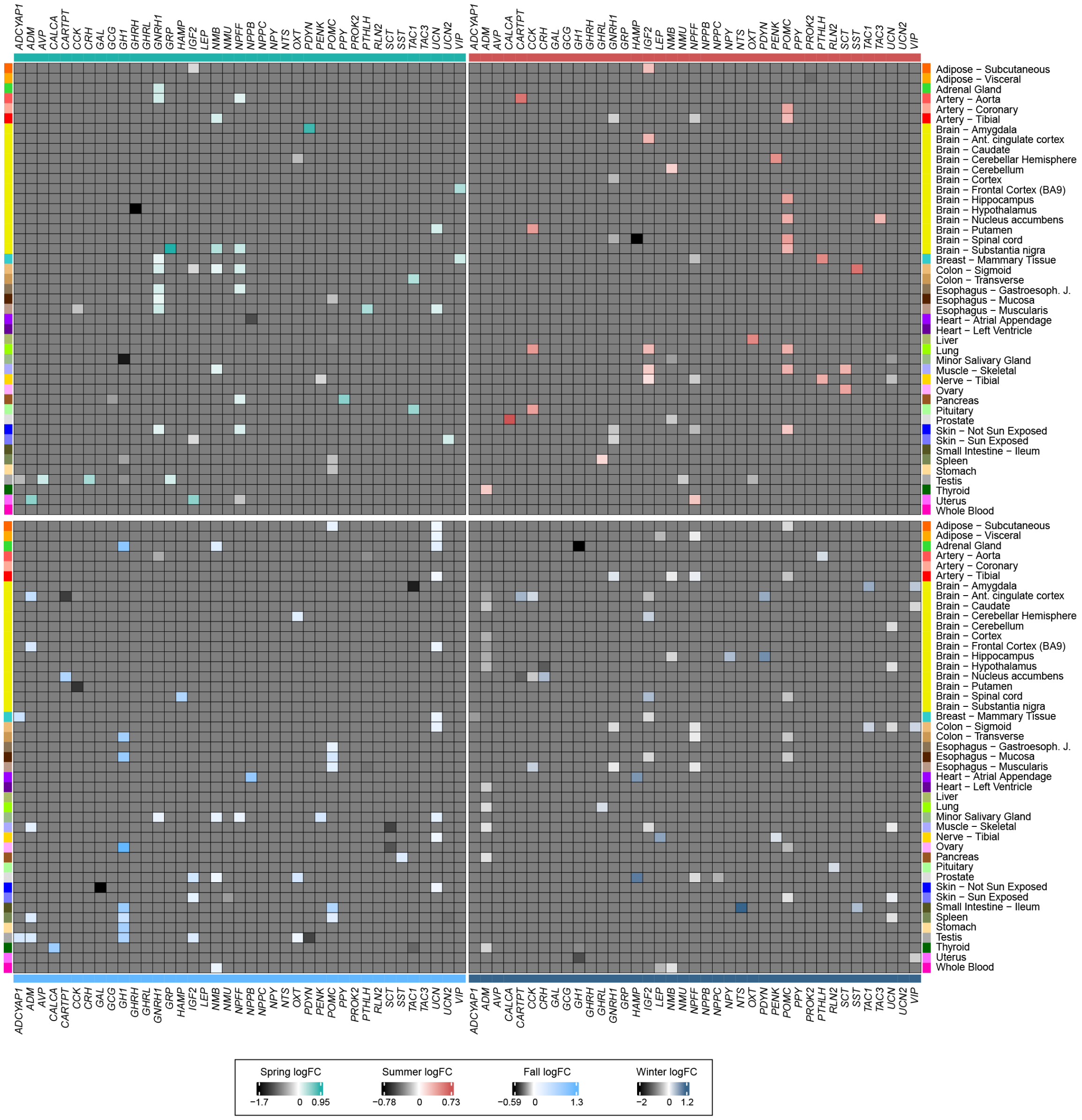


**Fig. S10 - Landscape of the seasonal variation of human hormone genes.** Seasonal *log₂* fold-change for the 39 hormone genes in the 45 tissues (all except vagina) with at least one hormonal gene showing seasonal expression.

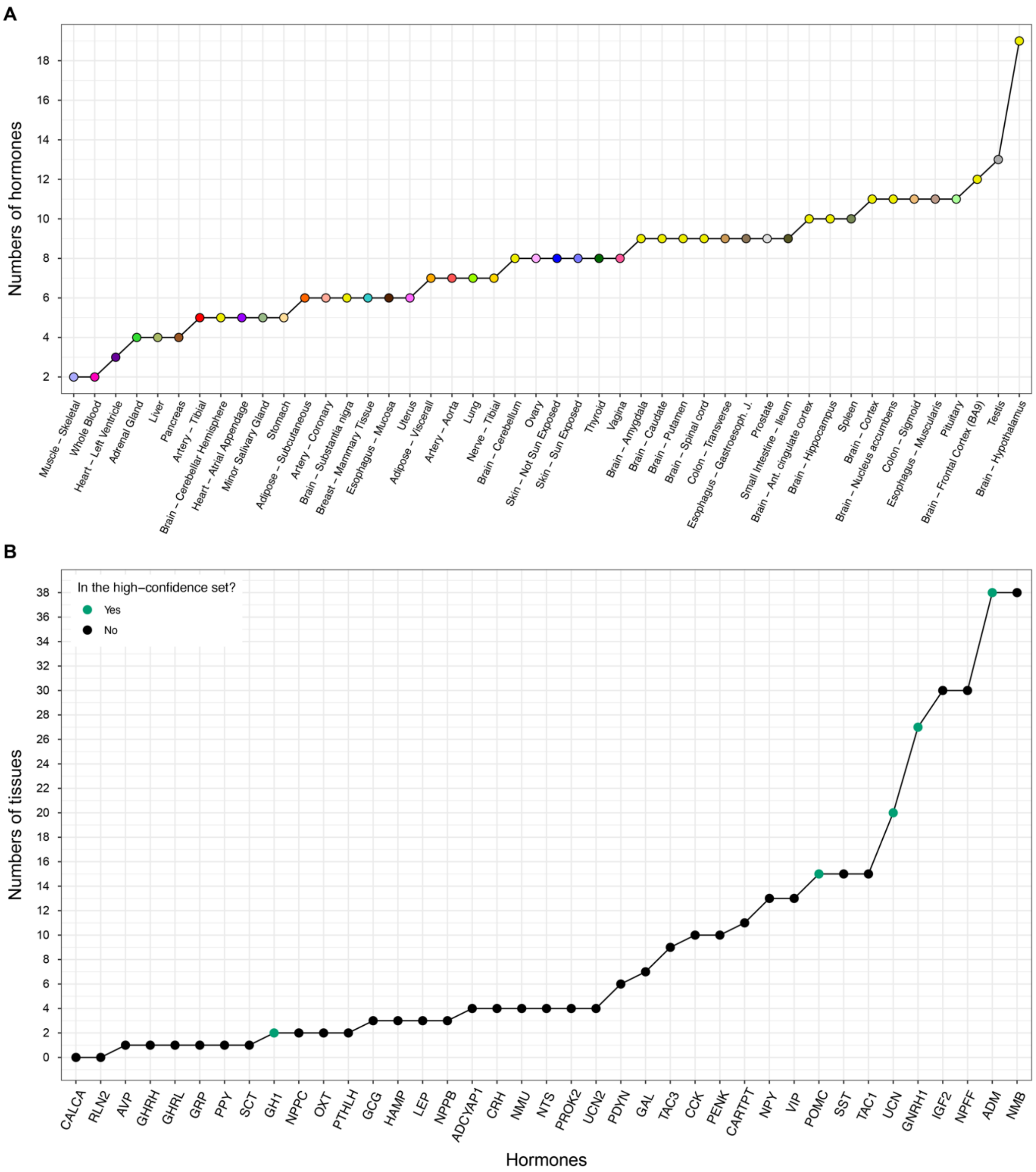


**Fig. S11 - Summary of the seasonal variation of human hormone genes.** (A) For each tissue, the number of seasonal hormone genes with a median expression TPM ≥ 5. (B) For each seasonal hormone gene, the number of tissues for which the gene had a median expression TPM ≥ 5.

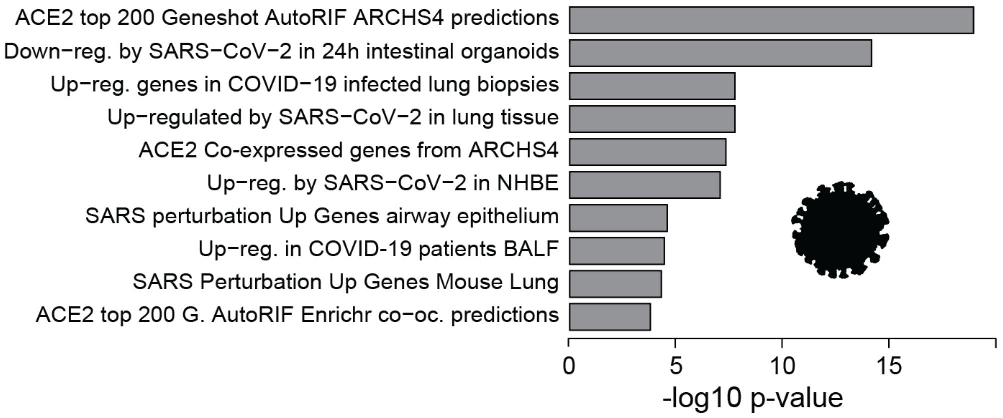


**Fig. S12 - Enrichment in COVID-19 related genes of the 192 strongly seasonal genes as computed by Enrichr.** *ACE2* Geneshot AutoRIF ARCHS4 predictions were obtained by Geneshot (Lachmann et al. 2019) combining genes previously published to be associated with *ACE2*, as well as genes predicted to be associated with ACE2 based on data integration from multiple sources, including co-expression matrices based on RNA-seq data and others (Kuleshov et al. 2020).

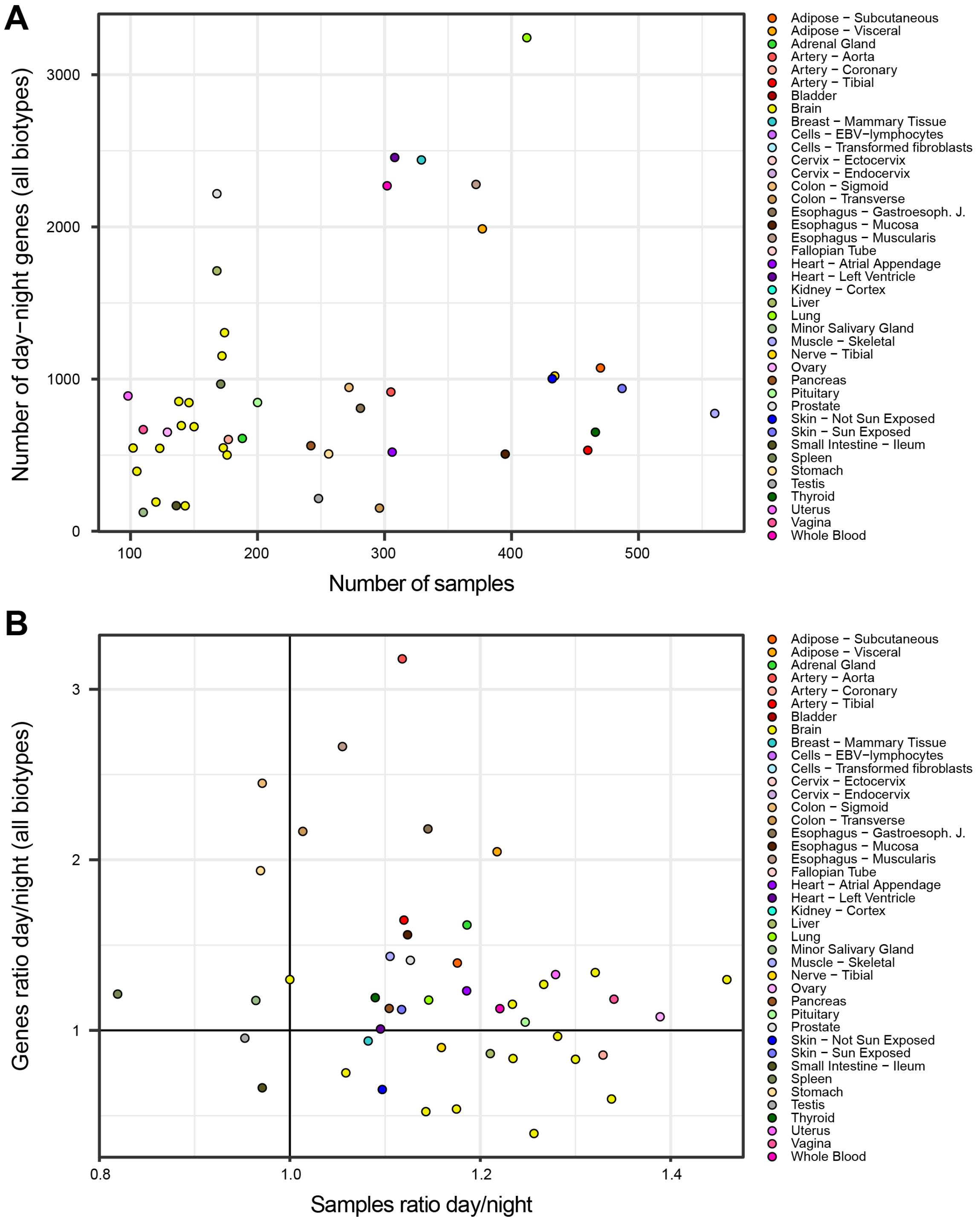


**Fig. S13 - Correlations between day-night variation and sample size.** (A) Number of day-night genes per number of samples. (B) Ratio of the number of day over the number of night genes vs the ratio of day over night samples.

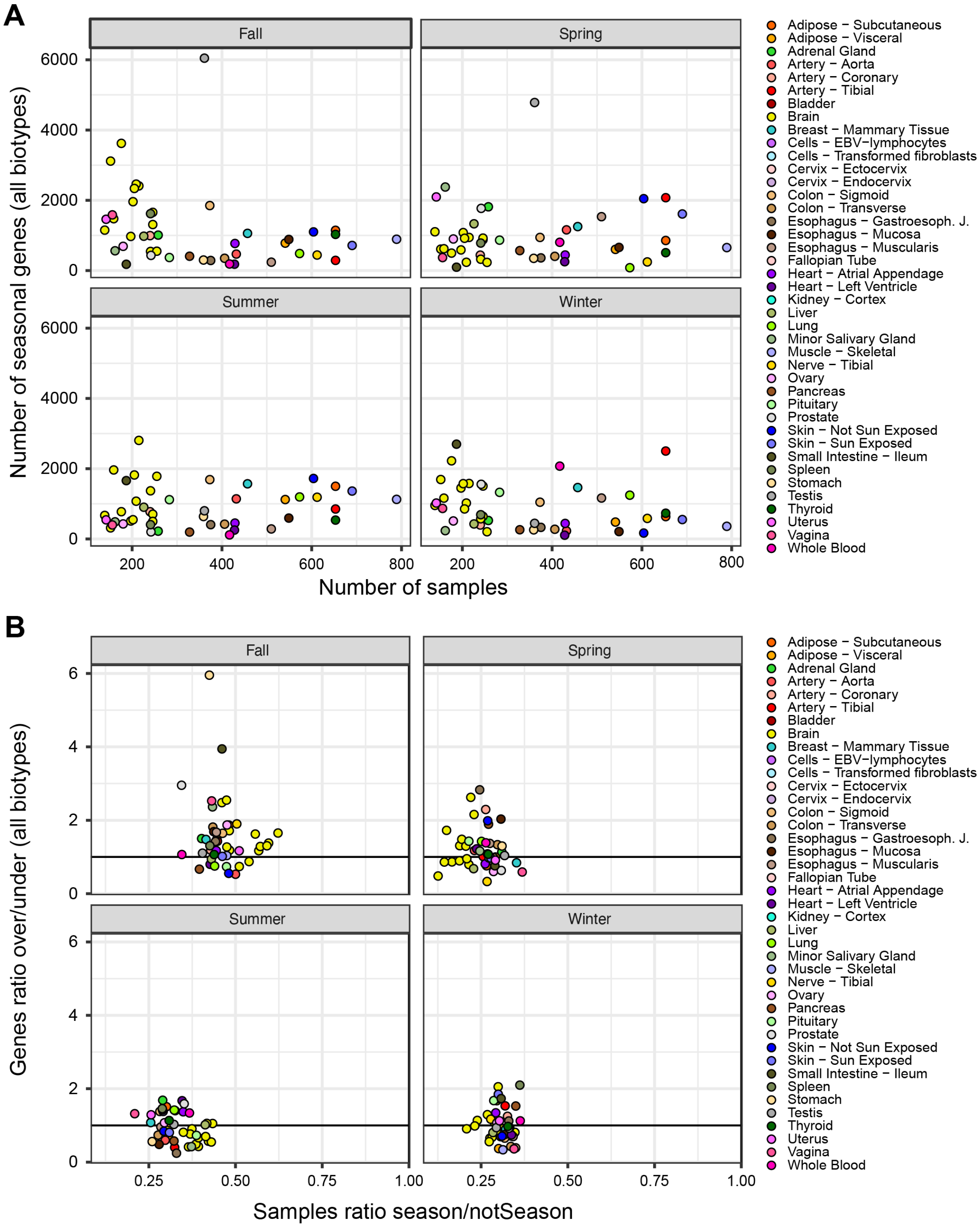


**Fig. S14 - Correlations between seasonal variation and sample size**. (A) Number of seasonal genes vs number of seasonal samples per season. (B) Ratio of the number of genes found over-expressed over genes found under-expressed per season vs the ratio of the samples from a specific season over all samples.
